# A single cell transcriptomic fingerprint of stressed premature, imbalanced differentiation of embryonic stem cells

**DOI:** 10.1101/2023.05.23.541952

**Authors:** Ximena Ruden, Aditi Singh, Teya Marben, Wen Tang, Awoniyi Awonuga, Douglas M. Ruden, Elizabeth Puscheck, Hao Feng, Daniel Rappolee

## Abstract

Cultured naïve pluripotent ESC differentiate into first lineage, XEN or second lineage, formative pluripotency. Hyperosmotic stress (sorbitol), like retinoic acid, decreases naive pluripotency and increases XEN in two ESC lines, as reported by bulk and scRNAseq, analyzed by UMAP. Sorbitol overrides pluripotency in two ESC lines as reported by bulk and scRNAseq, analyzed by UMAP. UMAP analyzed the effects of 5 stimuli - three stressed (200-300mM sorbitol with leukemia inhibitory factor +LIF) and two unstressed (+LIF, normal stemness-NS and -LIF, normal differentiation-ND). Sorbitol and RA decrease naive pluripotency and increase subpopulations of 2-cell embryo-like and XEN sub-lineages; primitive, parietal, and visceral endoderm (VE). Between the naïve pluripotency and primitive endoderm clusters is a stress-induced cluster with transient intermediate cells with higher LIF receptor signaling, with increased Stat3, Klf4, and Tbx3 expression. Sorbitol, like RA, also suppresses formative pluripotency, increasing lineage imbalance. Although bulk RNAseq and gene ontology group analyses suggest that stress induces head organizer and placental markers, scRNAseq reveals few cells. But VE and placental markers/cells were in adjacent clusters, like recent reports. UMAPs show that dose-dependent stress overrides stemness to force premature lineage imbalance. Hyperosmotic stress induces lineage imbalance, and other toxicological stresses, like drugs with RA, may cause lineage imbalance, resulting in miscarriages or birth defects.

## Introduction

During normal development, the placental and embryonic stem cells arise at the blastocyst stage before implantation into the uterus, and their proliferation accelerates from the blastocyst through gastrulation. Also, before and after implantation, stem cell differentiation to produce essential cell types occurs as the embryo prepares for gastrulation when the three germ cell layers are created and patterned (Puscheck et al., 2015; Puscheck et al., 2022). In general, stress homeostasis diminishes the growth trajectory of stem cells during implantation. Still, the embryo and its stem cells must meet developmental milestones for the development of parenchymal function necessary for cell survival and the organism’s survival. Specifically, sufficient 1^st^ differentiated lineage XEN must produce sufficient function despite stress-diminished growth observed in previous studies (Li et al., 2019; Slater et al., 2014). XEN functions include obtaining nutrition (Ross and Boroviak, 2020) and inducing head (Knoetgen et al., 1999) and heart (Madabhushi and Lacy, 2011) during gastrulation.

But most previous stress studies have focused on biochemical methods with small marker sets, with limited studies on changes in subpopulation sizes of initial naïve pluripotent stem cells. Previous studies examined naïve pluripotent stem cells using wild type and Rex1 promoter-red fluorescent protein (RFP) ESC (Li et al., 2016a; Li et al., 2016b), and the first essential differentiated parenchymal function of extra-embryonic endoderm (XEN) using Pdgfra promoter-green fluorescent reporter (GFP) ESC (Li et al., 2019) and Rex1-RFP ESC (Abdulhasan et al., 2021a). It is necessary to create XEN and its sub-lineages in order: first primitive endoderm (PrE), then structural parietal endoderm (PE), and nutritive visceral endoderm (VE) (Rappolee, 1999). A subpopulation of anterior visceral endoderm arises and is necessary to induce head organizer during gastrulation (Knoetgen et al., 1999; Madabhushi and Lacy, 2011; Zhou and Anderson, 2010). In mouse, this preparative period begins with naïve ESC formation in the E3.5-4.5 preimplantation inner cell mass. Then formative ESC function arises from E4.5 before implantation to E5.5-6.0 (Kalkan et al., 2017). Primed pluripotency peaks by the start of gastrulation at E6.5 and declines through gastrulation from E6.5-8.5 when germ cell layers arise and are patterned as the oct4-maintained pluripotency is lost.

Stress depletes ATP to meet homeostatic stress response while proceeding with the nearest emulation of normal development (Puscheck et al., 2015; Puscheck et al., 2022). Stress typically increases catabolism and limits the anabolism of many polymers, including proteins and mRNA. This leads to slower growth. The problem the organism needs to solve is the simultaneous successful cellular stress response with the developmental organismal response, so that interacting co-dependent tissues arise with sufficient parenchymal function from the initiation of stress to the milestone approximating each normal developmental event.

Our overarching hypothesis is that stress that diminishes anabolism and proliferation must force sufficient, timely differentiation, which includes a more focused adaptive strategy to cope with stress while providing sufficient differentiation from fewer-than-normal cells (Puscheck et al., 2015; Rappolee et al., 2012). Although previous studies use biochemical techniques and examine the average loss of naïve pluripotency and gain of 1^st^ lineage XEN, or FACS, to test for changes in sizes of these subpopulations, what is needed is a scRNAseq that provides information on integrated transcriptional programs that change during the ESC response to stress.

Although the focus here remains on hyperosmotic sorbitol and its dose-dependent effects informed by scRNAseq studies, all-trans-retinoic acid (ATRA or RA) was added as a normal morphogen for XEN and 2CEL (Iturbide et al., 2021; Iturbide et al., 2022) subpopulations arising from ESC culture. This enables comparison of stress-forced differentiation of XEN, despite LIF, with RA-directed differentiation during LIF removal. It should be noted that ATRA is also a recognized toxicant for pharyngeal development in later development but is a hypothetical toxicant earlier in the peri-implantation embryo (Huang et al., 2005a; Huang et al., 2005b; Huang et al., 2003; Rodriguez et al., 2006; Rutledge et al., 1994).

We report here, using bulk and scRNAseq, that stress suppresses transcript number and genes expressed per cell but increase premature XEN and its sub-lineages, primitive, parietal, and visceral endoderm with similar marker development but slightly less than RA in depth and breadth of marker induction for XEN or 2CEL. We report a “more from less” strategy of stressed ESC, where fewer cells, with less mRNA and fewer total genes, expressed, emulate the induction of XEN and its sub-lineages and the 2CEL ESC short sublineage in a way phenomenologically like RA.

## Materials and Methods

### Materials

Rex1-RFP ESC were created and used previously in toxicological HTS (Abdulhasan et al., 2021a; Li et al., 2016a; Li et al., 2016b). FUCCI muESC were a kind gift from Dr. Pierre Savatier (Coronado et al., 2013). ESC lines were tested for ploidy and mycoplasma contamination. DMEM was obtained from Thermofisher-Gibco. Glutamax and sodium pyruvate supplement solutions were from Life Technologies (Grand Island, NY). ESC-qualified EmbryoMax fetal bovine serum, 0.1% gelatin solution, and ESGRO^TM^ Mouse LIF medium supplement were from EMD Millipore (Billerica, MA). MEM non-essential amino acid solution, sorbitol, 2-mercaptoethanol, and other chemicals were from Sigma (St. Louis, MO). All-trans Retinoic Acid (ATRA/RA) and Sorbitol (Sor.) from Sigma (Cat# SLCB4143 and s3889, respectively). All reagents were used with protocols approved by the Wayne state university Institutional Biosafety committee.

## Methods

### Embryonic stem cell culture

Mouse embryonic stem cells (mESCs) were cultured as described previously (Li et al., 2016a; Li et al., 2019; Li et al., 2016b). FUCCI mESCs were cultured in the absence of feeder cells in FluroBright DMEM (Gibco, CAT#A18967-01) supplemented with 15% mESC-screened FBS, 2mM L-glutamine, 1mM sodium pyruvate, 1 mM nonessential amino acids, 0.1 mM 2-mercaptoethanol, and 1000 U/mL murine LIF on 0.1% gelatin-coated dishes at 37LJC in humidified air with 5% CO_2_ (Masui et al., 2008). Rex1-RFP mESCs were cultured similarly but with red DMEM without glutamine (Thermofisher #11960044) but supplemented with Glutamax instead. All ESC were optimized at passage for exponential growth during the stimulus period, which began ∼18hr after passage at 15% confluence onto 96-well plates. Rex1-RFP mESC were allowed to stay at normal stemness (NS), stimulated by removing stemness-maintaining LIF (normal differentiation/ ND), or stimulated with sorbitol positive control stress at 200mM, 250mM, or 300mM (S200, S250, S300). FUCCI mESC were also grown by NS, ND, and S300, with the additional stimulus: RA without LIF. Osmolality of ESC media with and without added 200-300mM sorbitol was determined previously (Slater et al., 2014).

### RNA isolation, cDNA library prep, and bulk RNAseq

FUCCI ESC grown under RA (-LIF), NS (+LIF, 0 dose stress), ND (-LIF control, 0 stress), and 300mM sorbitol stress (+LIF) conditions for 72hr were used to perform triplicated biological experiments to determine transcriptomic responses by bulk RNAseq. To this end, mRNA expression analysis was done by the Wayne State University Genome Sciences by the following methods. An aliquot of the RNA was assessed by microfluidics using the ScreenTape for the Agilent 2200 TapeStation. As done previously, the electrophoretogram, RNA Integrity Number (RIN), and the ratio of the 28S:18S RNA bands were optimized for the overall RNA quality. RNA-seq was used to determine expression profiles. Lexogen’s QuantSeq 3’mRNA-seq Library Prep Kit (FWD for Illumina) was used for building RNA-seq libraries from 0.1-200 ng of total RNA in 5 µl of nuclease-free ultrapure water. Libraries were quantified on the Qubit and Agilent 2200 Tapestation using the DNA High Sensitivity Screen tape. The barcoded libraries were multiplexed at equimolar concentrations and sequenced on an Illumina NovaSeq 6000. Data were demultiplexed using Illumina’s CASAVA 1.8.2 software. After quality was assessed (Andrews, 2010), and reads were aligned to the mouse genome (Build mm9) (Dobin et al., 2013) and tabulated for each gene region (Anders et al., 2015). Differential gene expression analysis was used to compare transcriptome changes between conditions (Robinson et al., 2010). Significantly altered genes (log fold change ≥ 2; FDR ≤ 0.05) were used to identify affected pathways (Huang et al., 2009).

To ascertain whether other stem cell lines besides FUCCI ESC had similar transcriptomic responses, additional data was analyzed from an exposure of Rex1-RFP ESC for 72hr using NS, ND, and three doses of sorbitol for dose-dependent effects: 200mM, 250mM, and 300mM (S200, S250, S300). These doses have been used in several previous studies in mouse ESC (Li et al., 2016a; Li et al., 2019; Slater et al., 2014), placental trophoblast stem cells (TSC) (Awonuga et al., 2011; Zhong et al., 2010; Zhong et al., 2007), and embryos (Xie et al., 2013; Xie et al., 2007; Zhong et al., 2010). This enables comparisons between the different models of dose-dependent changes in growth, expression of proteins, and phosphoproteins to correlate with transcriptomic changes. Transcriptomic assays of Rex1-RFP ESC were performed by similar methods and analyses as previously reported (Abdulhasan et al., 2021b).

### scRNA-seq data preprocessing and quality control

Using CellRanger 6.0.1 (Zheng et al., 2017)from 10x Genomics, raw Chromium scRNA-seq output was processed to align reads, generate feature–barcode matrices. The mm10 mouse reference genome was used to align the single-cell sequence reads. We specify the estimated number of cells to be 10,000 for each sample using the ‘`expect-cells=10000’`option to reflect the input cells further in cell extraction. The median number of genes detected per cell for five samples (NS, ND, S200, S250, S300) are 1,102, 547, 582, 625, and 561, respectively. Cell fraction reads are 28.2%, 24.1%, 22.6%, 27.1 %, and 20.4%, respectively. The median unique molecular identifier (UMI) counts per cell for each sample (NS, ND, S200, S250, S300) are 2,135, 818, 887, 93,9, and 800, respectively. In the quality control process using Seurat, we retained the genes that had expression in at least ten cells and cells that had an expression of at least 200 genes. Cells with more than 20% of mitochondrial sequencing reads were filtered out. After quality control, data normalization and scaling were (Hao et al., 2021a) performed in downstream analysis with Seurat 4.1.1 (Hao et al., 2021b; Stuart et al., 2019). All bulk RNAseq experiments were from triplicate independent biological experiments but scRNAseq came from single replicate biological experiments.

### Corroboration of select developmental, stress and loading control genes by qPCR

Total RNA was isolated using RNAeasy Mini Kit (Qiagen) and treated with DNase. The cDNA was prepared using QuantiTect Reverse Transcription Kit iScript (Qiagen) and assayed using SYBR Green by 7500 Fast Dx Real-Time PCR System (Applied Biosystems). Each biological experiment was performed independently three times and all genes were normalized to Rn18S rRNA as done previously (Slater et al., 2014; Yang et al., 2016). Relative mRNA expression levels were determined by the ddCT method (Livak and Schmittgen, 2001). Fold change of individual genes was determined by comparison to expression in cells cultured at 20% O2 potency conditions. Primers used are shown in **Supplemental Figure 4**. All primer pairs were checked for specificity using BLAST analysis and thermal dissociation curves to ensure amplification of a single product.

### scRNA-seq data clustering analysis (UMAP)

After preprocessing and data quality control, the objects of 5 samples (NS, ND, S200, S250, S300) were merged. We identified anchor genes to integrate five samples by using all 15,470 genes. To acquire the integrated object, we created a Seurat object list containing five samples (NS, ND, S200, S250, S300) and performed normalization as well as variable feature selection. Then a set of anchors between the Seurat objects was found. These anchors can later be used to integrate the objects using the ‘Intergration’ function. The objects were normalized using the ‘LogNormalize’ method, and the scale factor was set as 10,000. We found the integration anchors using the five objects and employed 1:30 dimensions using Canonical Correlation Analysis (CCA). After data integration, scaling, and principal component analysis were performed. The ‘FinderNeighbors’ function and the first main components were retained to construct a K-nearest neighbor (KNN) graph. We used the ‘FinderClusters’ function to cluster the cells with a resolution of 0.5. Eight clusters were identified and visualized using the UMAP dimensional reduction technique. Violin plots were generated by categorizing gene expression by their transcript levels in the eight different clusters.

Using the Wilcoxon rank-sum test, we used the ‘FindALLMarkers’ function in Seurat to identify the differential expression genes (DEGs) in each cluster. Adjusted P value and FDR were calculated. FDR□<□0.05 was defined as statistical significance. DEGs were defined as those genes that showed statistical significance. Once we clustered the cells, we identified the marker genes for each cluster. Cell type was then assigned and annotated to the clusters based on the identified marker genes.

### Statistical and Graphical Analyses

Data, except for the scRNAseq (N=1), are from at least three independent biological experiments and are presented as mean ±SEM (SD for **Fig.1). Fig.1**=two-way ANOVA Statistical analysis was done by student two-tailed *t*-test using GraphPad Prism 9.2.0 (GraphPad Software, San Diego, CA). Some graphs were initially composed and formatted using MS Excel (MS Office 365). And final formatting was done with Photoshop Elements Photoshop 13 Editor (Adobe systems incorporated, San Jose, CA). Venn Diagram< Ease David.

**Fig 1.**
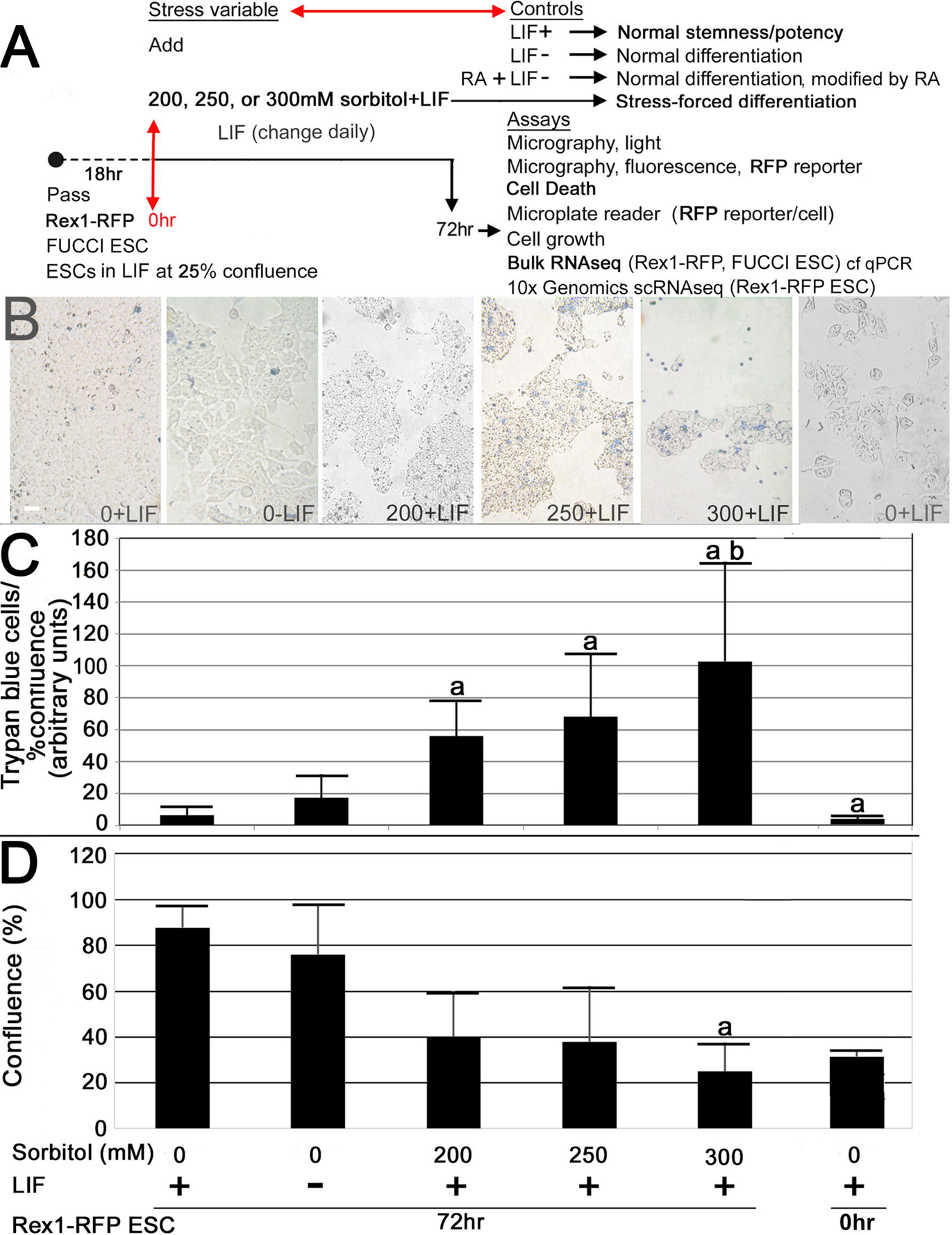
Rex1-RFP ESCs were stimulated to maintain NS (LIF+), undergo ND (LIF-) or stress-forced differentiation (LIF+; +200, 250, or 300mM sorbitol) (**<u>A, B</u>**), assayed for dose-dependent increase in number of trypan blue cells/epithelial confluence (**<u>C)</u>** or decrease in confluence (**<u>D)</u>**. (**<u>A)</u>** ESCs were exposed to media with LIF+ or LIF-with 200mM, 250mM or 300mM sorbitol for 72hr, micrographed (**<u>B</u>**) and ImageJ was used to quantitate fraction of confluence (**<u>D)</u>** and number of trypan blue cells/fraction of confluence (**<u>C</u>**). In B the white bar is 10um. The data shown are the X+-SEM from 3 biological replicates with (a) indicating significant difference from 72hr normal stemness (LIF+) and (b) indicating significant difference from ND (LIF-) (two-way ANO-VA; p <0.05).

## Results

### Fewer genes are expressed with stress, but there are more significant GO groups than NS, and a higher range of fold changes (FC) than ND, but few genes constitute the increased FC range

Rex1-RFP ESCs were cultured with increasing doses of sorbitol (S200, S250, and S300) or under stressless control conditions (NS/LIF+ and ND/LIF-) for 72hr (Time final-T_f_) and then assayed as shown in the experimental plan in **Fig.1A.** Stressed ESC-forced differentiated cells at 200-300mM were significantly different for trypan blue cells/% confluence compared with ESC in NS (**Fig.1B, C**). And ESC cultured 300mM, were significantly different from ND also (t-test, p<0.05) (**Fig.1C**) At Tfinal, ESC under NS and ND culture were ∼76-88% confluent, whereas ESC at 200-300mM sorbitol with LIF+ present were 25-40% (**Fig.1D**). Stressed ESC at 300mM were significantly decreased in confluence compared with ESCs during NS culture (p<0.05) (**Fig.1D**).

There is over a 3.2FC increase in the range of highest up- and down-regulated genes from ND to highest stress-forced differentiation (**Fig.2A**), but few genes undergo higher FC. Despite this large increase in the range of up- and down-regulated genes and a total of 15,562 genes expressed, the increased number of genes in the higher range of up- and down-regulated genes is only 12 and 10, respectively (**Fig.2A**). When 300mM sorbitol was compared with ND (**Fig.2B)**, there was a total 3.5FC increase in range of up- and down-regulated genes, but only 14 higher FC up- and down-regulated genes than ND. Surprisingly, only ∼28 genes were responsible for bringing out the maximum changes in the average fold change (28/15,562; ∼0.1%). Such a large effect brought out by such a small gene fraction may indicate that stress regulates an efficient response pathway.

**Fig 2.**
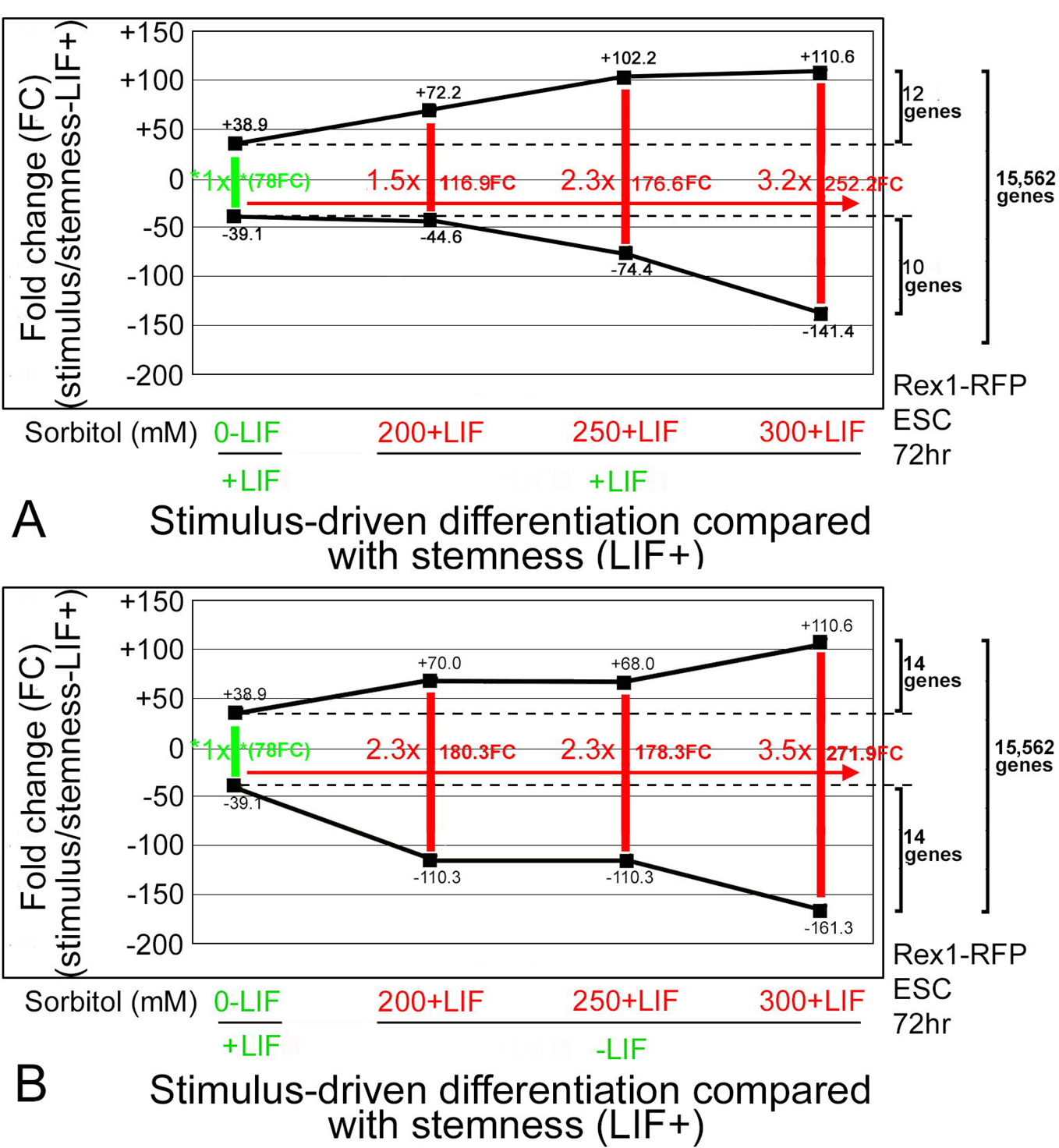
Stress forces dose-dependent increases in the highest up- and down-regulated genes, and a 3.2-3.5-fold change increase in range when compared with NS (**<u>A)</u>** or ND (**<u>B),</u>** but the up- and down-regulated genes at 300mM sorbitol is only 10-14 more than in highest regulated genes in ND. In triplicate experiments, Rex1-RFP ESCs were stimulated like Fig.1, lysed, mRNA isolated, checked for quality, reverse transcribed into cDNA, and counted by RNAseq. Stimulus-driven differentiation is either “normal” by LIF removal or “stress-forced” by adding sorbitol doses to ESCs despite LIF presence. In Excel spreadsheets, columns of FC between NS and ND and stress-forced differentiation compared with ND (**<u>A),</u>** or NS (**<u>B)</u>** were sorted from smallest to largest or vice versa. The highest up- and down-regulated gene was graphed, and the number of genes in the 300mM group above or below the highest normal upregulated or downregulated gene, respectively.

In contrast to higher FC at highest stress, the total number of expressed genes is lowest at the most elevated stress (**Supplemental Fig.1A).** And at every stratum of gene expression categorized by mRNA copy number/cell over four orders of magnitude (i.e., 10-10,000 highest expressed genes/cell), elevated stress at 250-300mM produces the lowest average transcripts per cell (**Supplemental Fig.1B-F).** Furthermore, both up- and down-regulated gene numbers are lowest at highest stress (**Supplemental Fig.2A-C**), but total, significant, and % significant GO groups are highest at highest stress (**Supplemental Fig.2D-F)**. The lower gene expression yet higher biological outcomes have been reported before (Judson et al., 2016), but this is a novel report and will be analyzed in the Discussion.

### For scRNAseq UMAP analysis, non-stress stimuli contribute more cells to clusters 0-1 and stress stimuli contribute more cells to clusters 2-7

To understand how dose-dependent stress (S200-S300 mM) and non-stress (ND/NS) stimuli induce cells with different differential gene expression (DGE), we analyzed the cells contributing to the UMAP by performing a clustering analysis. To understand stress and unstressed contributions to the clusters, a pie chart (**Fig.3A**) and its complementary graphed cell numbers and fractions (**Fig.3B**) were made. The five stimuli each contributed 10,000 cells to the UMAP to create eight clusters. If stress and non-stress stimuli were indistinct, there would have been an equal number of cells from each stimulus in each cluster, with 20% of the cells coming from each stimulus. Of the eight clusters, clusters 5, 6, and 7 produce outlying promontories and candidates for identifiable specific functional cell groups within heterogeneous culture (described in Fig.4-7) (**Fig.3C, D**). **Fig.3B** shows that distinct clusters (5, 6, 7) have the fewest cell numbers (∼400-1,600 cells), whereas clusters 0-4 have the most; in descending order from ∼16,000-5,000 cells. Clusters 2-7 are mostly stressed cells, as unstressed NS and ND cells make up less than a third of cells (**Fig 3A**). The clusters 5, 6, and 7 have the highest fraction of contribution, ∼75-85%, from the three stress stimuli (**Fig.3B, 3C**). The grouping of unstressed and stressed cells is shown when the UMAP is produced from the two stressless stimuli, ND, and NS, or all three or only the two highest hyperosmotic stress doses **(Fig.3E**) S200 turns on the maximum number of genes (**Fig.1, supplemental fig.1**).

**Fig 3.**
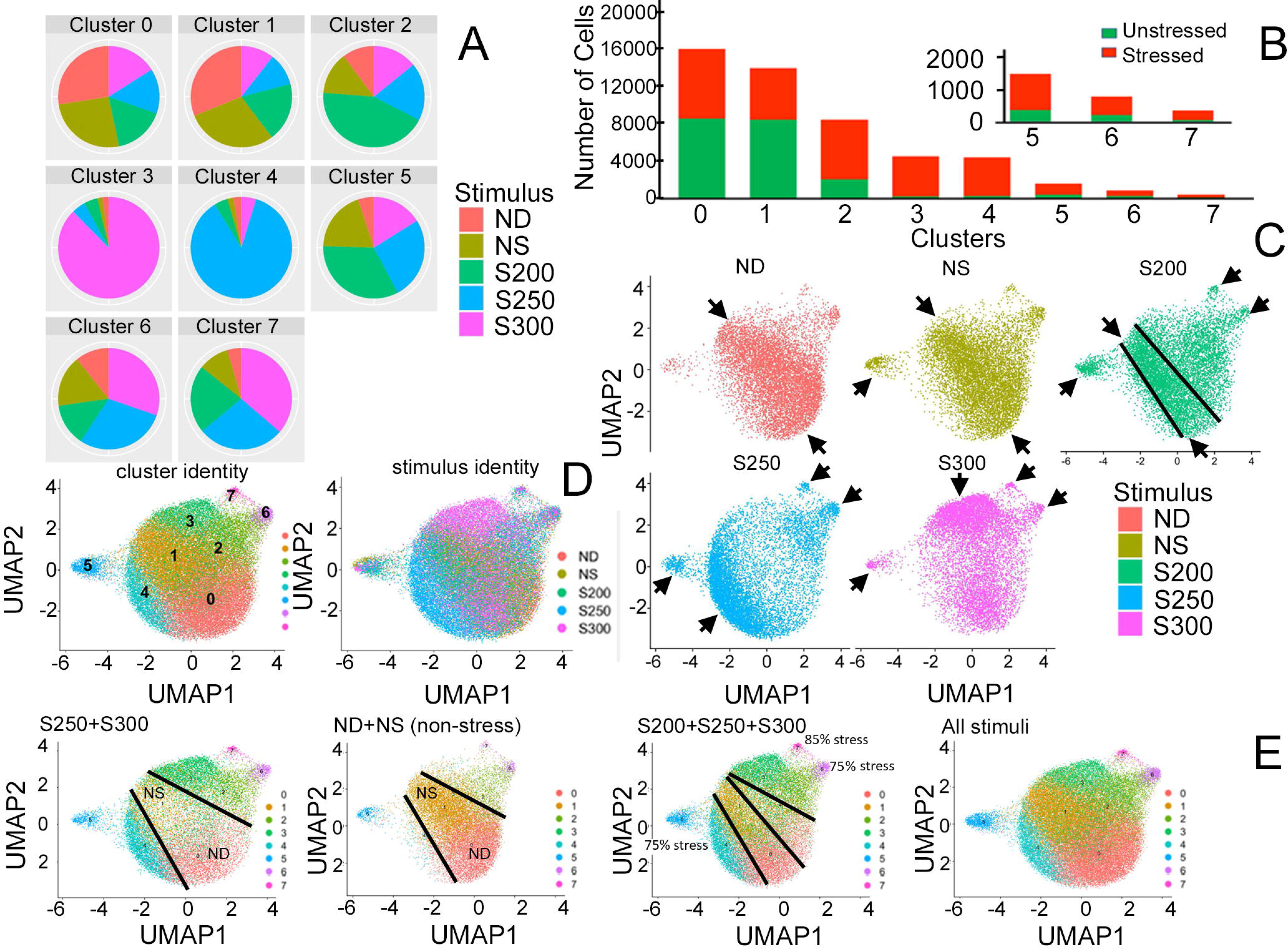
General stress dose dependent DGE edginess compared with non-stressful NS and ND centrality. (**A**) shows the fraction of cells contributed to each UMAP cluster by stress from 200-300mM sorbitol, and non-stress, ND and NS, and the cell numbers contributed to each cluster by each stimulus (**<u>B</u>**) graph shows the cell number in each cluster from 5 different stimuli **(<u>C)</u>**shows UMAP central cell contribution (indicated by black arrows) of non-stress ND and NS, and highest UMAP edge contribution by highest stress doses 250mM and 300mM sorbitol, and high UMAP edge contribution by 200mM sorbitol with some central area contribution (compared central 200mM area defined by black lines <u>in **Fig.D and Fig,E**</u> with the low donation of 250mM and 300mM sorbitol in **<u>Fig.E</u>**) to cluster 0 and 1, primarily populated by NS and ND. **(<u>D)</u>** compares the composite stimulus-dependent cell contribution on the right to the 8 clusters mapped to the UMAP on the left. **(<u>E)</u>**shows the contribution to UMAP edges by different stress doses and the central cell contribution by stressless ND and NS.

**Fig 4.**
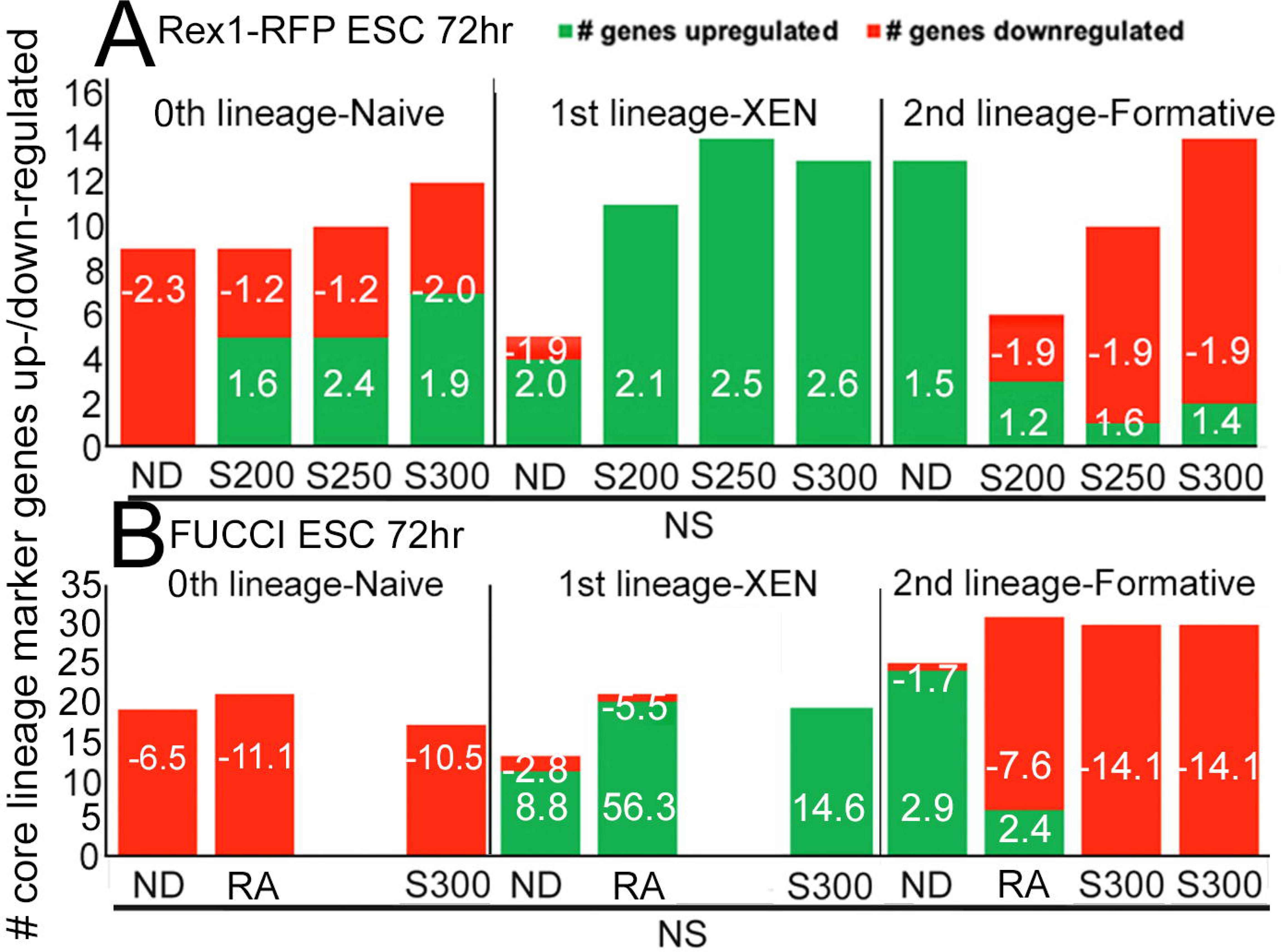
Histogram shows effect of 3 stimuli on the core markers for the first three ESC lineages compared with initial NS culture for Rex1-RFP ESC (A) and FUCCI ESC (B). The X-axis shows the five stimuli (ND, RA, and sorbitol doses; S200, S250, S300); the y-axis indicates the number of genes being upregulated (in green) or downregulated (in red) for those lineages. The data labels in the graph shows the average FC for the upregulated/downregulated genes.

### Bulk RNAseq shows ND, RA, and sorbitol decrease core naïve gene transcripts and increase core XEN gene transcripts. RA and sorbitol suppress but ND increases formative pluripotent gene transcripts

We tested for override of NS by stress by analyzing bulk RNAseq, comparing NS with the highest dose of sorbitol (S300), or lower sorbitol doses (S200-S250), ND and All-Trans Retinoic acid (RA) after 72hr exposure (FUCCI ESC). RA, S200-S300, and ND cause the cells to downregulate the genes for 0^th^ lineage/naïve, indicating that the three stimuli decrease Naïve pluripotency in Rex1-RFP ESC (**Fig.4A**) or FUCCI ESC (**Fig.4B**). All three stimuli caused the ESC to upregulate 1^st^ lineage/XEN genes. For the 2^nd^ lineage/formative pluripotency, ND was shown to upregulate the genes of the formative stage, whereas S200-S300 and RA were shown to downregulate the genes for the formative pluripotency and thus suppressing the 2nd lineage.

The two ESC lines are similar in increasing XEN and decreasing Formative pluripotent cells for hyper-osmotic and RA stimuli and increasing Formative pluripotent cells for ND. RA is a stronger stimulus in both average FC for up- and down-regulated genes, and for the total number of significantly-changing genes. The two ESC lines have greatest differences and are more variable in the strength of stimulus-dependent departure from naïve pluripotency.

### scRNAseq confirms that sorbitol induces premature lineage imbalance to 2CEL and XEN sub-lineages PrE, PE, and VE compared with ND

Bulk RNAseq analysis reveals increased XEN as a biochemical average. A UMAP was used to investigate whether stress induced all three XEN sub-lineages. ICM/naïve pluripotent cells delaminate first into primitive endoderm (PrE), then these cells contact the basal surface of the trophectoderm and become migratory parietal endoderm (PE). PrE then becomes visceral endoderm (VE) which later develops into the anterior visceral endoderm (AVE). The top 3 marker genes for each UMAP cluster are shown in bubble plots **(supplemental fig.3A**), and for clusters 6 and 7 (**Fig.5A and 5B**). Col4a1 and Col4a2, PrE/PE marker genes, were highly expressed in cluster 7, (**Fig.5A**). And Krt8 and Krt18, VE marker genes, were detected in cluster 6 (**Fig.5B**). Col4a1/2 markers and Krt8/18 marker pairs are in the top three highest expressed and with the highest subpopulation size, for clusters 7 and 6 respectively, and are markers of the PrE/PE and VE XEN subpopulations markers (Despin-Guitard et al., 2022; Illingworth et al., 2016; Martinez-Ceballos et al., 2005; Wang et al., 2002). This suggests that cluster 7 is PrE/PE and cluster 6 is VE. Cells expressing PE markers. LamC1 and LamB1 (Li et al., 2004; Murray and Edgar, 2001; Zhuang and Gudas, 2008), were assayed and co-detected in cluster 7 (**Fig.5C**). PEG10 and H19, markers of many cells but co-expressed in XEN, were tested for co-expression, and detected in cluster 6 (**Fig.5D**). Similarly, SPARC and Lama1 genes, although expressed in many cell types, are co-expressed during the transition of PrE to PE, and co-expressing cells were detected in cluster 7 (**Fig.5E**). Thus, cluster 7 contains the early lineages of XEN-PrE and PE and Cluster 6 has later VE sub-lineage of XEN.

**Fig 5.**
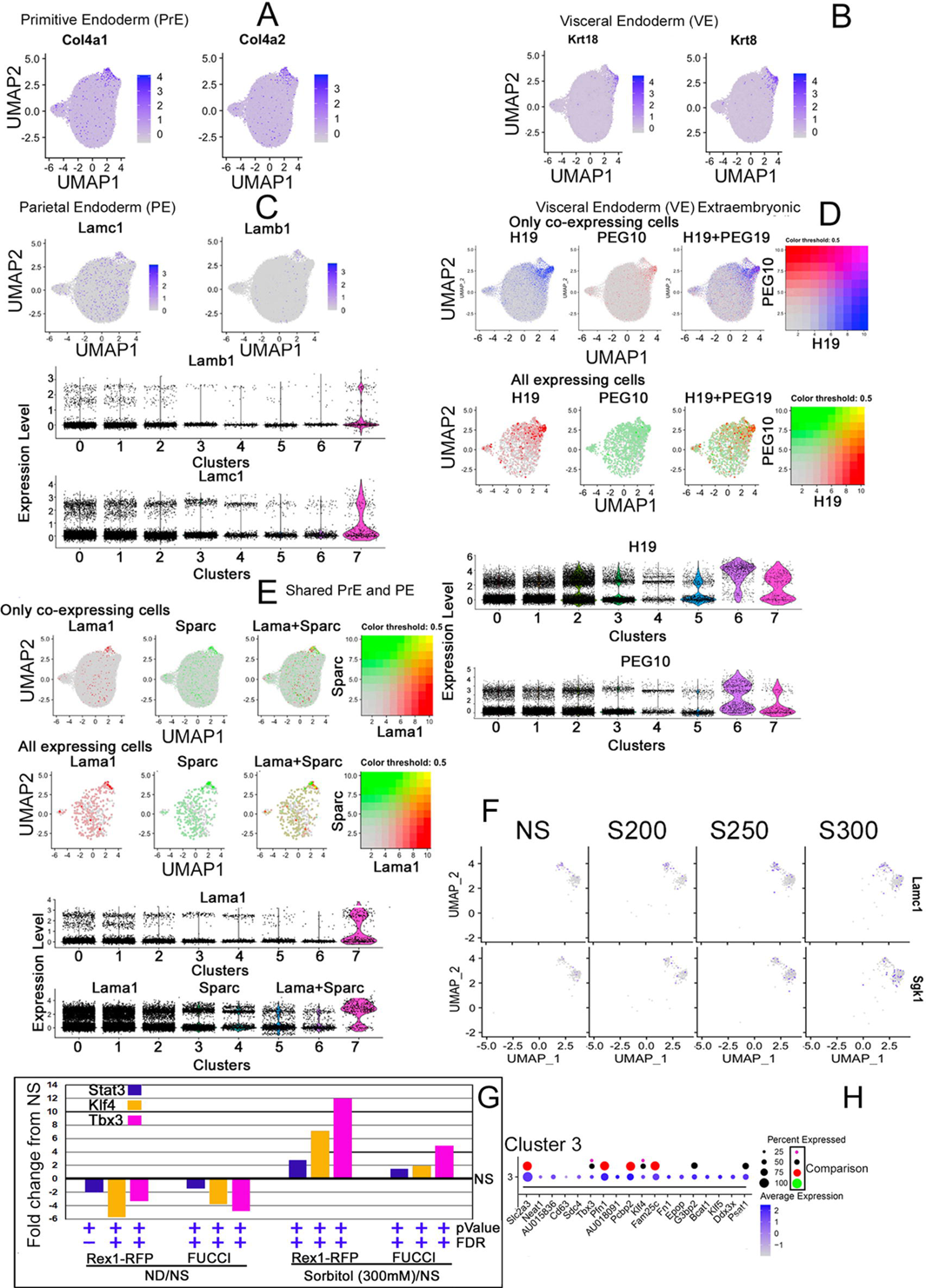
DGE analysis shows that cluster 7 contains PrE and PE, and cluster 6 contains VE gene markers sub-lineages induced by stress-forced override of naïve stemness. **(<u>A</u>**) Shows XEN subpopulation PrE gene markers localized to cluster 7. **(*<u>B</u>***) Shows XEN subpopulation VE gene markers localized to cluster 6. (**<u>C</u>**) Shows XEN subpopulation PE. (**<u>D</u>**) Shows co-expression of two XEN subpopulation markers of VE in 6 and spreading into cluster 2. (**<u>E</u>**) Shows co-expression of two markers shared during the transition of PrE into PE in cluster 7. (**<u>F)</u>** Dose-dependent increase of the marker genes for XEN sub-lineages with increasing stress. (**<u>G)</u>** Shows analysis of bulk RNAseq 72hr exposures of Rex1-RFP or FUCCI ESC and FC induced by ND vs sorbitol 300mM FC from initial NS culture (+ and – show pValue and FDR significance). Stat3, Klf4 and Tbx3 are standard LIF receptor response mRNA. (**<u>H)</u>** Shows 18 highest bubble plot expression levels and % cells expressing for cells in cluster 3. Comparison reference bubble sizes were replicated, and color coded for 25, 50, 75 and 100% expressing cells in cluster 3 and shown above standard bubbles to reference sizes. Note Klf4 and Tbx3 increase are in highest 18 expressed genes in cluster 3, which is 85% sorbitol 300mM sourced cells.

We hypothesized that sorbitol induces XEN in a dose-dependent manner. **Fig.5F** shows the cells expressing markers genes for XEN, and the number of cells increases in a dose-dependent manner although dose-dependence is not statistically significant by UMAP analysis.

**Fig.5G,H** show two aspects of expression occurring in cells largely induced by sorbitol 300mM in cluster 3. **Fig.5G** shows that three genes downstream of the LIF receptor are downregulated 72hr after LIF removal. Stat3, Klf4, and Tbx3 are significantly downregulated in Rex1-RFP and FUCCI-ESC during ND and upregulated by S300 (**Fig.5H)** and top 18 bubble plot genes Klf4 and Tbx3 are highly expressed for % cells (25-50%) and expression levels (1.5-2/scale of 2). Together the data suggest that cluster 3 has 300mM sorbitol stress-induced gene expression of genes downstream of the LIF receptor, at a higher level than NS.

To confirm the quantitation of gene expression by bulk RNAseq, we also tested the NS, NS, and sorbitol 200-300mM sorbitol by qPCR via the ddCt method (**Supplemental figure, 4, 5**). Loading controls Gapdh and 18S were mostly within 2-20% of each other for all stimuli and altogether only two stimuli created an expression above 20% different from the other stimuli, both for Gapdh. 18S was the more stable loading control. For the other developmental and non-developmental genes, all comparisons were not significantly different for bulk RNAseq and qPCR.

The UMAP shows 2CEL, typically reported in NS culture or after RA induction, is strongly induced by sorbitol (**Fig.6**). In **Fig.6A**, the presence of 2CEL markers is shown in cluster 5 of the UMAP. RA is known to induce 2CEL and sorbitol was similar. UMAPs were shown to have cells highly expressing Platr31 and GM21761 in cluster 5, suggesting that cluster 5 is 2CEL. Fbx015 and Nelfa, markers of many cells but known to be co-expressed for induction and function in 2-cell embryo or in the 2CEL subpopulation in ESC (Hu et al., 2020; Tokuzawa et al., 2003; Wang et al., 2021b), were tested for co-expression, and found in cluster 5 (**Fig.6B and 6C**). We test the hypothesis in **Fig, 6D** that stress causes a dose-dependent increase in the 2CEL sub-population. UMAPs were created for marker genes for 2CEL-Platr31 and Fbxo15, for increasing sorbitol doses (NS<S200<S250<S300), and the expression of 2CEL markers increased with increasing stress, supporting our hypothesis.

**Fig 6:**
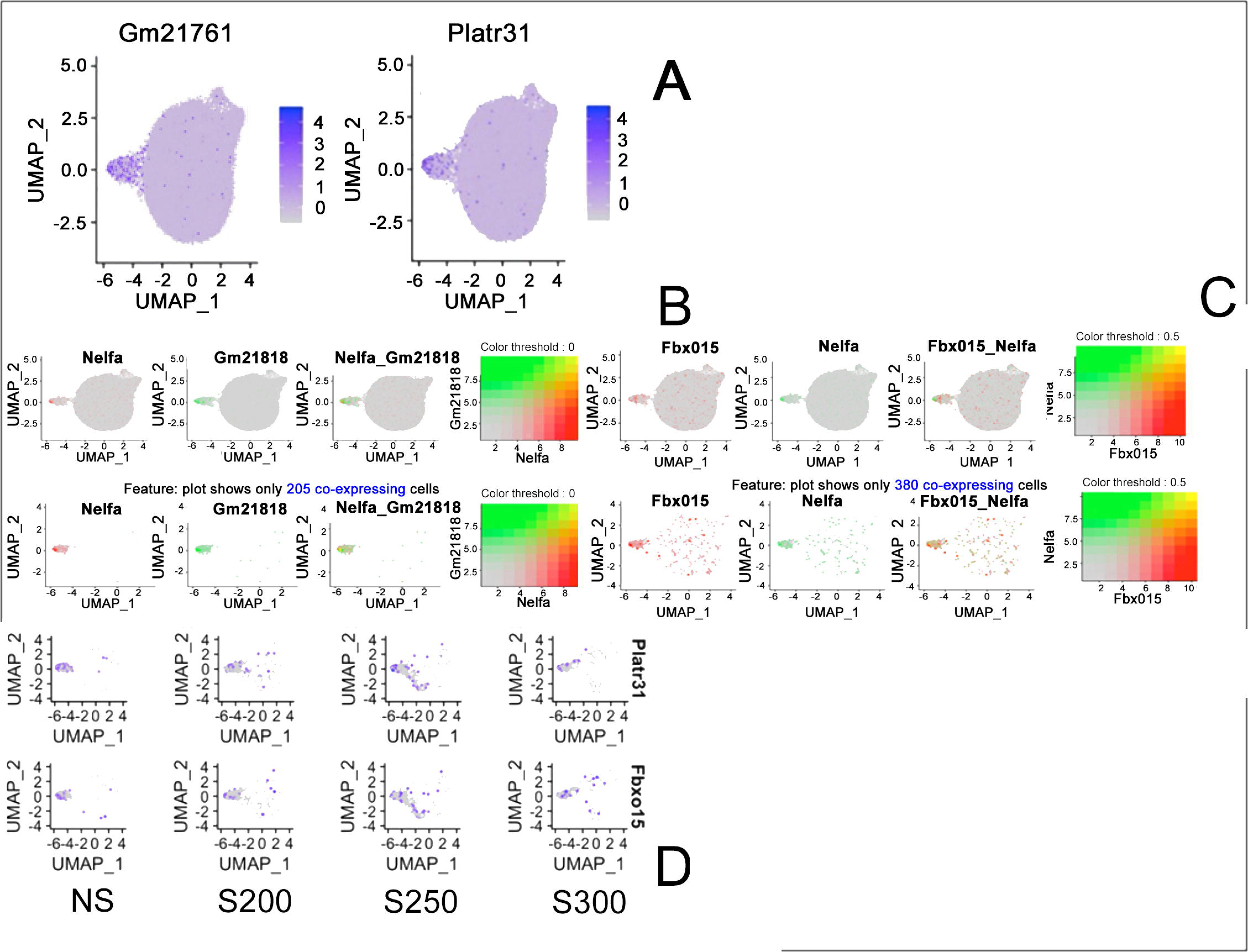
**DGE analysis shows that cluster 5 contains 2-cell embryo-like cells.** (**<u>A)</u>** shows 2CEL gene markers localized to cluster 5. **<u>(B and C)</u>** shows the co-expression of two 2CEL markers localized to cluster 5. (**<u>D)</u>** Shows dose-dependent decrease (Platr31) and dose-dependent increase (Fbx015) in marker genes for 2CEL.

### Formative markers arose, but not significantly by 72hr of LIF removal

Formative markers were induced by ND but suppressed by sorbitol (and RA) by bulk RNAseq (**Fig.4**). Heat-maps showed ND up-regulated the core formative pluripotent genes in Rex1-RFP and FUCCI ESC (**Fig. 7A)** equivalent to XEN at E5.5 (**Fig.7B).** So scRNA-seq was used to test for induction of formative pluripotent cells. **Fig.7C** shows that most cells from the unstressed stimuli (ND/NS) are found in the UMAP central part. The genes expressed by the highest cell number with the highest expression from the bubble plots (**Supplemental Fig.3)** for cluster 0 were L1td1, Ncl, and Atrx. These genes are known to regulate embryonic stem cell functions and may be involved with formative pluripotency (Emani et al., 2015; Iwabuchi et al., 2011; Percharde et al., 2018; Wong et al., 2010). Expression of Otx2 (known head-organizer gene)(Boroviak et al., 2018; Kondoh, 2018; Neagu et al., 2020; Yang et al., 2014) and Dnmt3a/3b (Betto et al., 2021; Boroviak et al., 2015; Kinoshita et al., 2021), both methylation enzymes that are up-regulated, enabling formative pluripotency, were also analyzed. Cells expressing these genes were found but not strongly localized to cluster 0 (**Fig.7D and E**). There was some tendency to be expressed more than in other clusters. Co-expression of formative pluripotency markers was stronger in cluster 0 (**Fig.7F**), and there was a tendency to higher co-expression. Expression and co-expression of formative marker genes in clusters 0 versus 1 are expressed by higher number of cells in cluster 0 than in cluster 1 (**Fig.7G**) but at a much lower multiplier (1.5fold) than pairs of XEN genes in cluster 6 (100fold) and 7 (200fold) compared with other clusters. Thus, formative sub-transcriptomic programs are expressed in cluster 0>1. We hypothesized that stress stimuli were suppressing formative pluripotency. We found more unstressed cells expressing the formative genes, and the average expression for the formative genes is higher in unstressed versus stressed cells. (**Fig.7H**), which supported our hypothesis that stress (Sorbitol) suppresses formative lineage.

**Fig 7.**
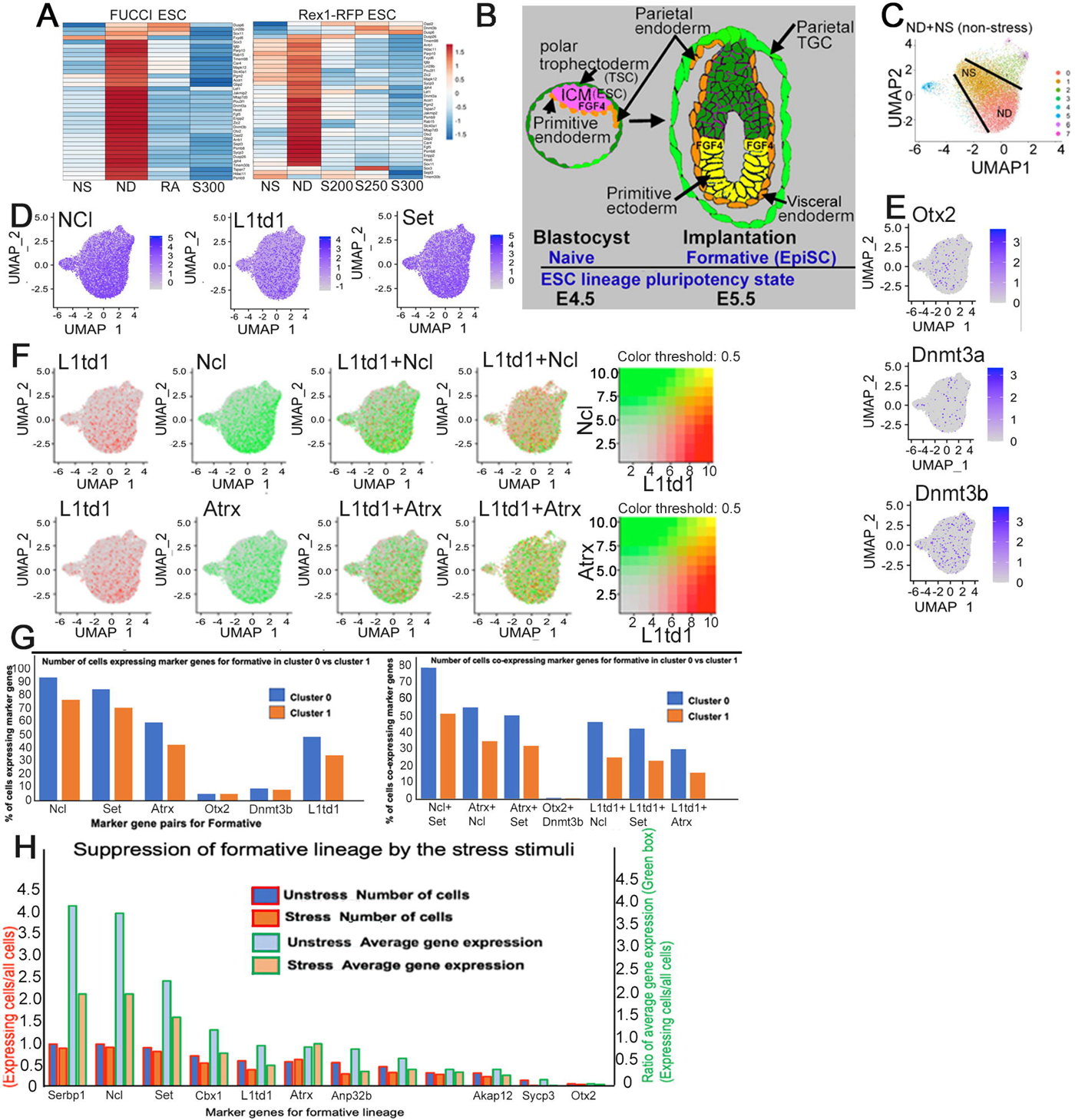
DGE analysis shows that cluster 0 contains a formative lineage sub-transcriptomic program. **(A)** Heat-maps show genes up/down (red/blue) regulated by the stimuli (bottom) for core formative lineage genes in FUCCI and Rex1-RFP ESCs. **(<u>B</u>)** ESC in their naïve pluripotency stage E4.5 (pink) is shown on left and ESCs in their formative pluripotency stage E5.5 (yellow cells) are shown on the right. **<u>(C</u>)** A UMAP shows the center is made up of NS and ND, unstressed cells. (**<u>D and E</u>**) UMAPs illustrate the cells expressing the formative marker genes localized to cluster 0. (**F**) UMAPs show the cluster-locations of cells expressing marker genes (columns 1-2). Column 3 shows a merge of the two genes on the UMAP, resulting in gradient change. Column 4 shows all the cells co-expressing the two marker genes, and their varying expression level is shown by different colors, as indicated in column 5. **(<u>G)</u>** Figures show the fraction of cells expressing (left) and co-expressing (<u>right)</u> the formative lineage marker genes in cluster 0 versus cluster 1. The y-axis shows the fraction of cells and genes are shown on the x-axis. **(<u>H)</u>** The graph shows the suppression of formative lineage by stress stimuli. Two y-axes are shown: the left (red) y-axis shows the ratio of the number of cells expressing marker genes (red outlined histogram bars) and the right (green) y-axis shows the ratio of average expression of marker genes (green outlined histogram bars). The X-axis shows the formative lineage marker genes. Dark and light blue bars show unstressed cells, and dark and light red bars indicate stressed cells.

Although formative marker genes were expressed in cluster 0 more than other clusters, the formative sub-transcriptomic program did not form a separate cluster. We tested the strength formative pluripotency from ND compared with 16hr formative than the 25hr formative marker expression reported previously (Kalkan et al., 2017). ND by LIF removal for 72hr was not like Kalkan’s 16hr or 25hr (data not shown). Formative might have needed more time to arise more completely. The results from Bulk RNAseq analysis could predict the formative lineage that would eventually arise with extended culture.

### Placental genes are up-regulated by stress in a dose-dependent manner

Bulk RNAseq reveals upregulation of placental genes as a biochemical average. The key placental genes are higher expressed with stress>ND, and stress>NS (**Fig. 8A**). This suggests that stress might be turning the ESC into placental cells. We further analyzed with a larger marker gene set for placental genes and the results from bulk RNAseq on FUCCI ESC showed that ND mostly down-regulates placental genes (**Fig 8B**) while RA and S300 mostly up-regulated the placental genes (**Fig. 8C and 8D**). Bulk RNAseq for placental genes was also performed on Rex1-RFP ESC to determine the induction of placental genes with stress in a dose-dependent manner. ND did not induce placental marker genes (**Fig. 8E**), corroborating our findings with FUCCI ESC. S200 upregulated placental genes (**Fig. 8F**), S250 and S300 upregulation of placental marker genes was progressively higher (**Fig. 8G, N**). The ICM does not make placenta and hence it was expected to see no up-regulation of placental genes in ND. We used scRNAseq to test whether placental cells arose and found cells expressing the marker genes for placental genes on the UMAP, but co-expression was low (**Fig 8I**). However, co-expression for placental genes was still found to be 4.7 times more in cluster 6 and 7, than any other cluster, indicating the presence of placental marker genes in stressed clusters. Interestingly, Hand1 and H19 placental markers are two of the three highest expressed and highest represented cell subpopulations in cluster 2 and H19, Krt8 and Krt18 are the three highest expressed and highest represented cell subpopulations in VE of adjacent cluster 6 (bubble plots in **Supplemental Fig.3**). This is interesting as a recent study using UMAPs and lineage tracing showed that normal visceral yolk sac and trophoblast from mouse conceptuses are together the most distant from embryonic ectoderm but similar to each other (Chan et al., 2019). In contrast, here the predominant cell types in cluster 2 and 6 are 70-75% induced from stress override of normal stemness.

**Fig 8.**
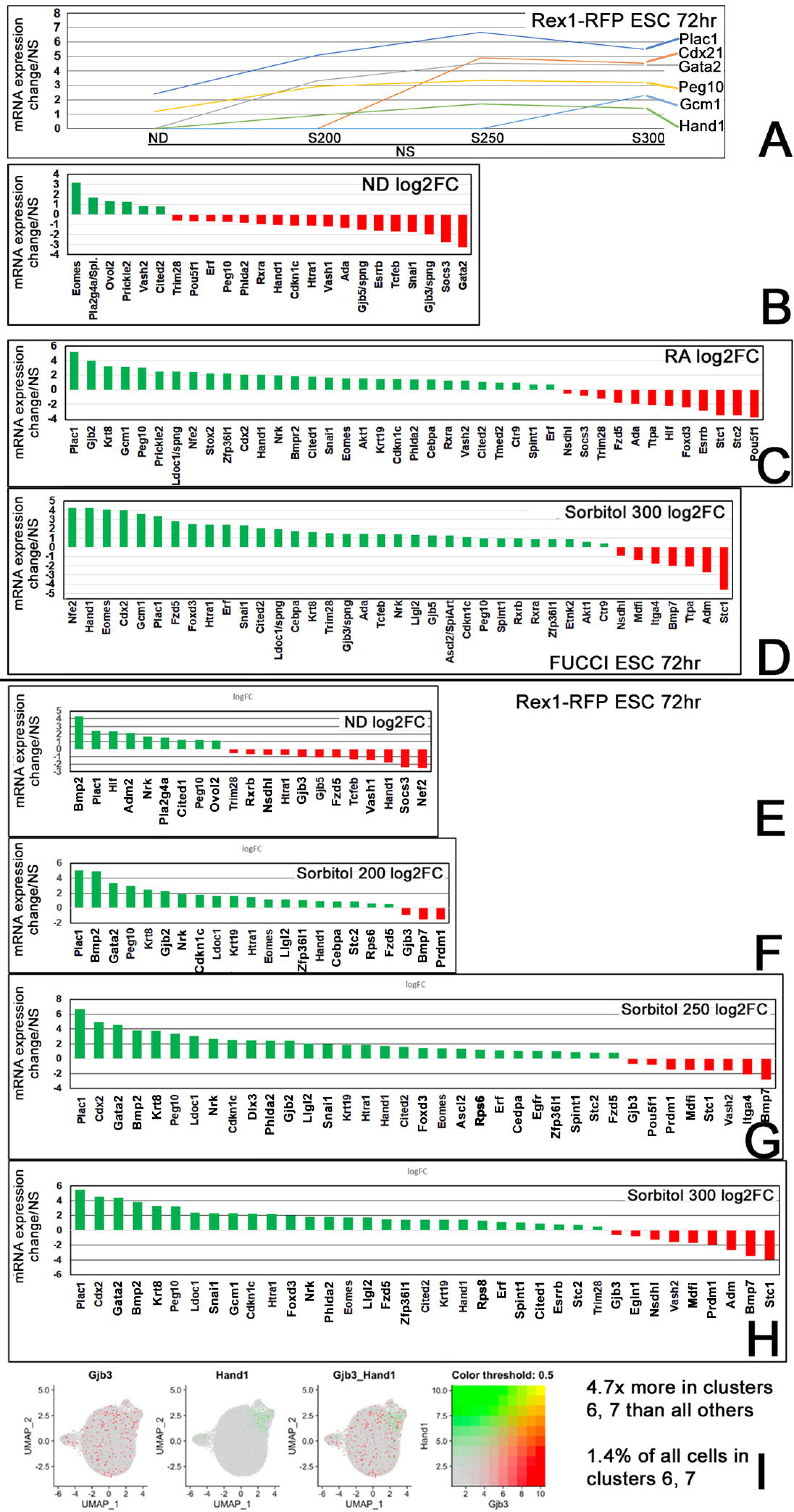
Placental markers increase from NS to ND to stress-forced override **(A)**, and generally increase with RA and sorbitol, but not ND **(B-H).** However, scRNAseq does not detect significant numbers of placental cells **(I)**. **(A)** shows gene expression for marker placental genes on the right for the stimuli listed on the x-axis. **(B-D)** show data for FUCCI ESC and **(E-H)** for Rex1-RFP ESC. Marker placental genes on the x-axis are up/down (green/red) regulated for ND **(B, E)**, RA **(C)**, S200 **(F)**, S250 **(G)**, and S300 **(D, H). (I)** The 2 UMAPs on the left show the cells that express key marker placental genes. The third from the left shows a UMAP with a merge of the two genes, resulting in gradient change; their varying expression level is shown by different colors, on the right. <u>[All genes shown are p-value significant genes, error bars not shown]</u>

### Rex1-RFP and FUCCI-GFP ESC agree for reported XEN and 2CEL induction

A comprehensive list of significantly-induced XEN genes induced in Rex1-RFP ESC by S200-S300 and S300 and RA induced in FUCCI ESC was identified (**Fig.9**). We hypothesized that induction of 1^st^ lineage/XEN is more dire than 2CEL. To test the hypothesis, we took RA XEN data (Iturbide et al., 2021; Iturbide et al., 2022) for time-dependent induction. We compared the 2CEL and XEN genes induction in our data from FUCCI ESC analysis with previous reports (Han et al., 2018; Mohammed et al., 2017) and saw that 2CEL had highest overlap with the mid (early-arising and late-persisting) whereas XEN overlapped with late-arising and late-persisting time (**Fig.10A and 10B**). This stress induces 2CEL in ESC culture before XEN; refuting our hypothesis and suggesting 2CEL induction is more dire.

**Fig 9.**
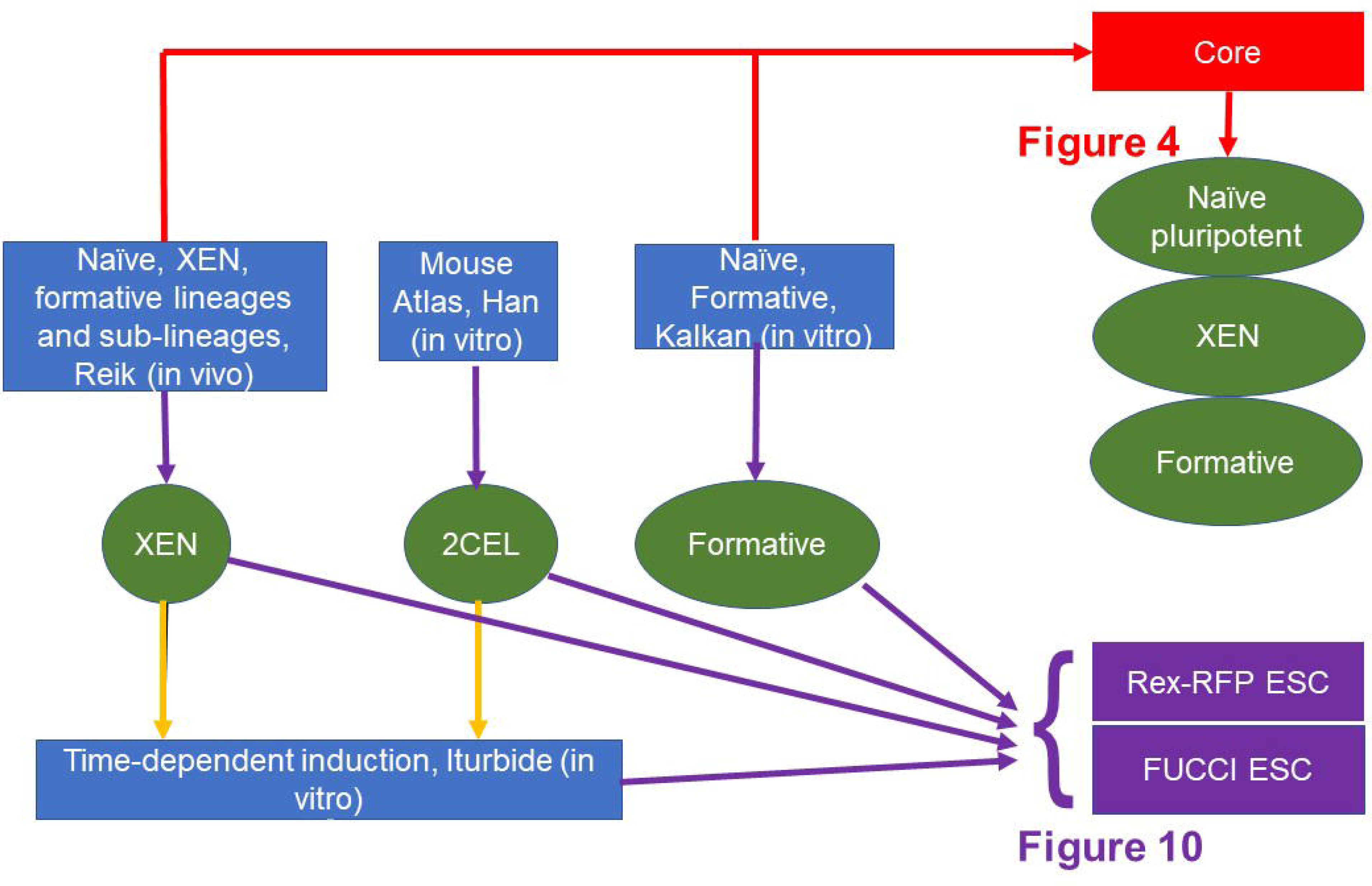
Figure showing different comparisons of published literature with data obtained from Rex1-RFP and FUCCI ESC. Markers for in vivo induction of XEN, it’s sublineages and formative lineages from Mohammed et a 2017, were compared with in vitro time-dependent induction of XEN and 2CEL from Iturbide et al 2021. Han et al 2017 was used for 2CEL markers and compared with Iturbide. Red lines show various data that were cross-referenced to obtain the core genes for naïve, XEN and formative used in Fig.4. Purple lines indicate the cross-referencing of various datasets against FUCCI and Rex1-RFP ESC to analyze the data shown in **Fig. 10.**

Past RA studies document 2CEL and XEN induction in ESC, but hyperosmotic stress is less defined. Bulk RNAseq data in **Fig.4B**, show that RA and S300 similarly induce 2CEL and XEN lineages. However, S300 is weaker than RA, as indicated by the magnitude of gene expression (e.g., insets of the bar graph). RA and S300 are similar but RA. We hypothesized that the induction of 2CEL and XEN is similar to RA and S300. To test this hypothesis, we compared the expression profiles of RA and S300 from FUCCI ESC with previous time-dependent expression profiles (Iturbide et al., 2021; Iturbide et al., 2022). RA and S300 share a similar percentage of genes across early-arising (no genes shared), early-arising and late-persisting (7% and 4% of shared genes), and late arising and late persisting (32% and 33% shared genes) (**Fig 10B**). These results support our hypothesis that the timing of induction of 2CEL and XEN by RA here and previously is similar.

**Fig 10.**
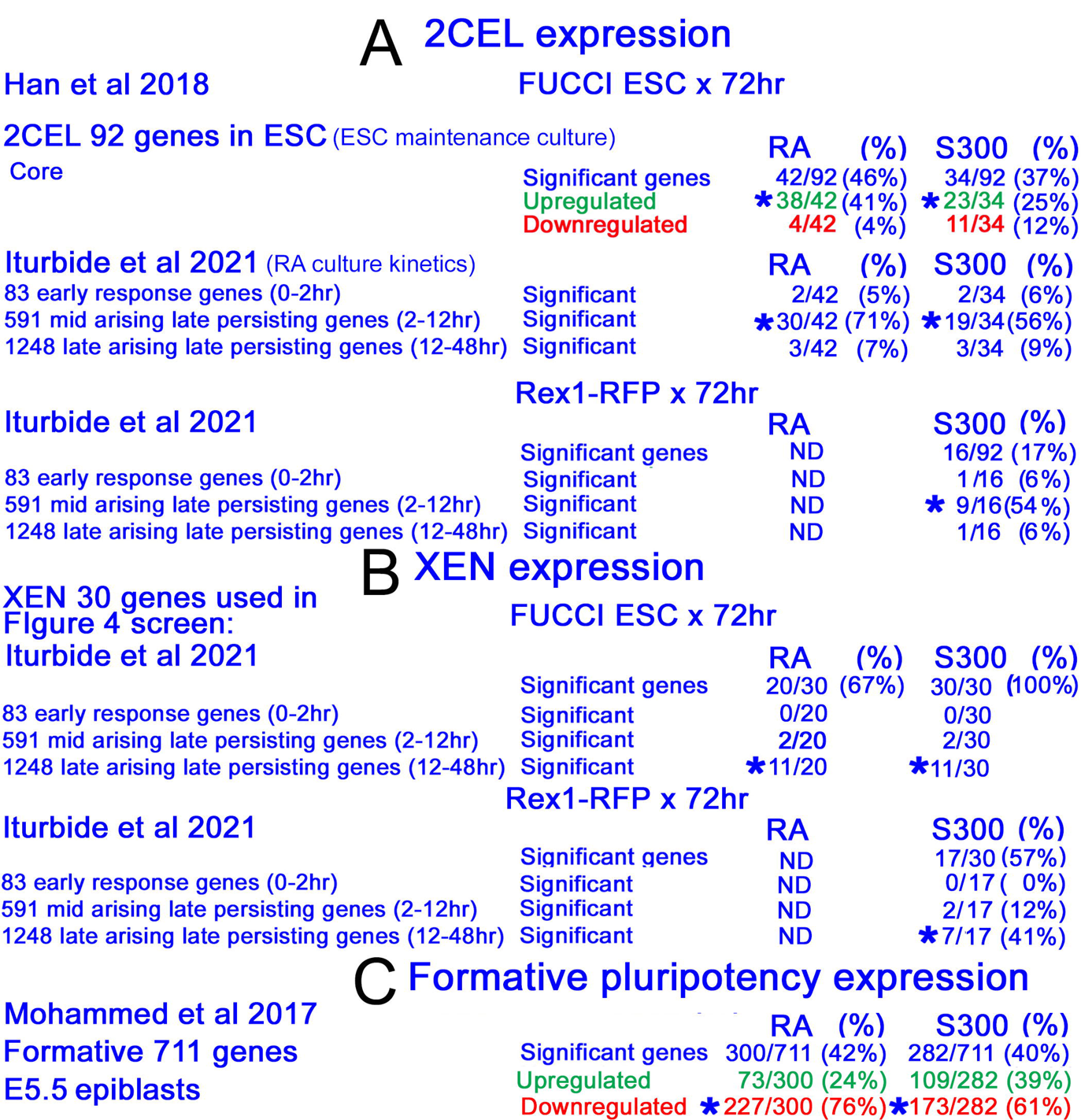
Figure showing data from various cross-referencing shown in Fig.9. The papers used for comparisons are shown on the left and the number of genes shared for FUCCI and Rex1-RFP are shown on the right. The asterisks indicate the group having highest fraction of genes shared. (**A**) shows comparisons of 2CEL, (**B**) shows comparisons for XEN and (**C**) shows comparisons for Formative lineages.

We hypothesized that in vivo induction of XEN and 2CEL would be similar to in vitro induction. We compared genes for PrE for days E4.5, 5.5, and 6.5 in vivo (Mohammed et al., 2017), with the early-arising, early-arising and late-persisting, and late-arising and late-persisting XEN and 2CEL in vitro (Iturbide et al., 2021; Iturbide et al., 2022)(**Fig 10B**). There are 3 time-points in vivo and in vitro from both studies. We hypothesized that the genes from E4.5 would have the most shared genes with early arising, genes from E5.5 would have the most shared genes with early arising and late persisting, and E6.5 would have most genes shared with late arising, and the late persisting group. We observed that for all days, E4.5, E5.5, and E6.5, most genes were shared or best identified with the late arising and late persisting group. One tendency is that early arising in vivo are later arising in vitro. This could mean that early arising in vivo genes do not arise with RA induction until late. It is probable that culture delays development, which is why there is so much overlap with late-raising and late-persisting groups.

From the scRNAseq for formative pluripotency (**Fig.7**), we showed that RA and S300 were similar in the suppression of formative lineages. To confirm this, we compared the list of formative genes from E5.5 in vivo studies (Mohammed et al., 2017) with FUCCI ESC markers here and found that RA and S300 suppressed the formative lineages (**Fig. 10C**). This supports our hypothesis that RA and S300 are similar in suppression of formative lineages and also shows that in vitro and in vivo studies report similar findings.

## Discussion

Clustering analyses of scRNA-seq data on the effects of five stimuli on cultured ESC produced one cluster with cells having 2CEL properties and two clusters having properties of XEN and its three XEN sub-lineages. Further marker analysis showed that PrE and PE were in cluster 7 and VE was in cluster 6. Previous reports of hyperosmotic stress and RA increasing XEN used only small marker sets (Li et al., 2019; Slater et al., 2014). It was confirmed that RA and hyperosmotic stress have similar effects in inducing 2CEL and XEN transcripts, but that the response to RA usually has higher FC and involves a larger number of genes. But RA modified ND during LIF removal and hyperosmotic stress overrode stemness in the presence of LIF.

Interestingly, between cluster 0, which had markers of unstressed NS cells, and cluster 7, which had the markers of initial PrE and PE sub-lineages, was cluster 3. Cluster 3, like cluster 7, had cells that were 85% derived from stress stimuli, but had elevated levels of gene expression for LIF receptor; Stat3, Klf4 and Tbx3. The data suggests that stress induces higher levels of LIF receptor response than NS itself. This seem strange, but is similar to previous reports that the highest levels of Oct4 protein occurs transiently in the earliest PrE adjacent to the ICM (Palmieri et al., 1994). Oct4 at higher levels is known to anticipate XEN induction (Niwa et al., 2000), and Oct4 is required cell-autonomously in the pre-PrE cells in the blastocyst (Frum et al., 2013). These data suggest that elevated Oct4 or other elements of LIF R activation may be part of a transient state between naïve pluripotency and PrE during normal or stressed differentiation.

Increased XEN and decreased ESC growth are hallmarks of stress responses of cultured ESC. It is interesting that ultrasound studies or early post-implantation human embryos show that when yolk sac to embryo size ratios increase, reporting a similar lineage imbalance in vivo, this associates with miscarriage (Detti et al., 2020a; Detti et al., 2020b). These data suggest a similarity in lineage imbalance of in the embryo in vivo, and similar effects in cultured ESC may predict miscarriage.

### The lack of GO groups for early lineages creates unique challenges and solutions in interpreting DGE data

GO groups are used to understand omics programmatic changes but have low fidelity with the analyses of early lineages of ESC. No GO groups exist for formative, primed pluripotency, or XEN lineages of early ESC lineage. The DGE analysis of bulk RNAseq data suggests GO groups identifying bone and kidney development in cultured ESC differentiated by LIF removal. But short-term LIF removal does not induce germ layer markers at gastrulation that would enable kidney cells and makes GO group analysis less credible for assigning developmental functions.

Since almost all genes in GO groups change significantly, these genes can be used to refine searches of gene expression, cross-referencing published bulk and scRNAseq data. Placental GO group genes showed that many genes were upregulated by stress more than normal development. But scRNAseq analysis did not show increased placental cell number. Human embryonic stem cells reprogrammed to the naïve state, can transdifferentiate into the TSC lineage (Dong et al., 2020; Dong and Theunissen, 2022), and thus it is possible that mouse ESC may do the same with the correct stimuli. In addition, adjacent clusters 2 and 6 have interesting properties of majority of cells induced by stress. By bubble plot analysis, in cluster 2, placental markers Hand1 and H19 are expressed in highest fraction of cells at highest levels, and in cluster 2 VE markers H19, Krt8, and Krt18 are expressed. Similar to this, a recent lineage tracing study placed in conceptuses place VE and placental UMAP clusters adjacent and distantly to embryonic body and tail cells (Chan et al., 2019). However, that study was in unstressed cells derived from the normal mouse conceptus, and studies here stress overrides stemness to drive cultured ESC into VE cells with an adjacent cluster where cells have possible placental function or future fate.

Similarity of DGE transcriptomic changes previously reported for RA were found to be like hyperosmotic stress override of LIF-induced naïve stemness here. Since RA induces naïve pluripotent ESC transition to 2CEL and XEN cells (Iturbide et al., 2021; Iturbide et al., 2022; Rodriguez-Terrones et al., 2018), we compare this with hyperosmotic override of LIF and naïve pluripotency. RA addition with LIF removal, or sorbitol override of LIF here, 2CEL markers were more like “early arising, later persisting” 2CEL arising before 12hr and persisting through 48hr reported previously (Iturbide et al., 2021; Iturbide et al., 2022). In contrast, RA or sorbitol induced XEN markers from bulk RNAseq, which were found to be like late-arising, late-persisting. Thus, the stress-induced need for 2CEL gene expression is more “dire” than XEN here because 2CEL is early-arising (before 12hr in the previous RA time course) and XEN is late-arising (after 12hr) DGE induced by hyperosmotic stress and RA. Or it could mean that developing XEN requires a multi-step transcriptional sequence that requires longer than inducing 2CEL.

Naïve pluripotency varies with different ESC culture protocols. Traditional culture media using serum with LIF shows that LIF removal induces 2CEL gene expression (Kalkan and Smith, 2014) and in studies here with serum, LIF removal with RA, or sorbitol override despite LIF, induces 2CEL. The ability to obtain less heterogenous pluripotency at Naïve/ground state using 2i+LIF in chemically defined media (Marks et al., 2012), 5i+LIF(Theunissen et al., 2014) or larger inhibitor sets higher inhibitor screens (Khan et al., 2021) using the distal oct4/ICM/naïve promoter reporter ESC. However, 2i+LIF does not enable 2CEL induction. Serum allows for a more heterogeneous maintenance culture and stress response.

Over 2,000 DGE markers are expressed in 2-cell embryos, but only 92 DGE markers are expressed in the transient, ∼1-2% 2CEL (Han et al., 2018). Hyperosmotic stress overrides naïve pluripotency to induce 2CEL of ∼1-2% subpopulation size in the UMAP cluster 5, but in bulk RNAseq sorbitol does not induce Zscan4c, but RA does. A comparison of RA and sorbitol induced Zscan4 levels here was compared to four levels of Zscan4 associated with transient subpopulations in RA-induced 2CEL (Rodriguez-Terrones et al., 2018), and similar to other comparisons RA induced mostly mid-to-high Zscan-associated genes, and the sorbitol was weaker inducing mostly low-to mid Zscan-associated genes (data not shown) and did not induce Zscan4 like RA.

### Reservations about current models of stress exposure in vitro

Media for Formative pluripotency/ EpiLC ESC have been formulated (Wang et al., 2021a). It is not known if any single medium enables the flexible equal capability to differentiate to all three germ layers at gastrulation, which is needed for unbiased testing of imbalanced differentiation for toxicological purposes. scRNAseq showed that 2i+LIF produces cells resembling ICM, and EpiSC media emulated the epiblast from embryos at the start of gastrulation, but DGE for ESC in EpiLC culture were most different from E5.5 epiblasts measured ex vivo (Posfai et al., 2021). Additionally, EpiLC and EpiSC may be usable for maintenance of formative and primed pluripotency and for testing for stress override, but the capability of “ND” control to emulate all later differentiation events is not clear.

### XEN induction is robust in override of stemness to increase mRNA and protein markers from several reports

Differences in clusters 6 and 7 were robust because they confirm similar subpopulations size, FC, and specific DGE changes reported previously. For example, the subpopulation of about 1% XEN in clusters 6 and 7 here reflects the same subpopulation size from the same dose and duration of sorbitol that was previously reported after FACS analysis of Pdgfra-GFP ESC (Li et al., 2019). Dab2 and Lrp2 XEN markers upregulated by hyperosmotic stress assessed by qPCR (Slater et al., 2014) and Dab2 protein by immunoblot (Li et al., 2019), had a very similar fold change to 10fold higher number of XEN genes reported here from bulk RNAseq. Induction of XEN in override of stemness is quantitatively reproducible.

Majority unstressed cells in clusters 0 and 1 trended from naive toward formative pluripotency. In vivo, naïve pluripotent cells don’t exist 48hr after they form, and by that time, naïve have become formative and trending toward primed pluripotency. In vitro, heterogeneous naïve pluripotency supported by serum and LIF persisted for 72hr after LIF removal. The transition from naïve to formative is incomplete and closer to 16hr reversible formative than the 25hr irreversible formative reporter previously (Kalkan et al., 2017). The transition to formative is weak and could be accelerated with less heterogeneity by use of 2i+LIF.

### Strategies for cross-referencing results here with previous reports

Stress at the highest levels induced more cells on UMAP perimeters, whereas non-stressful ND and NS are generally more central in cell distribution on the UMAP. If the stress did not induce any effect, each cluster would consist of 6,225 cells (50,000 cells would have contributed equally to make the 8 clusters, and the fraction of non-stress cells would be 40% (coming from NS and ND), and stress cells would be 60% (coming from three sorbitol doses)). The results indicated that most cells (∼30,000) were found in clusters 0 and 1, where most cells were unstressed. Most of the stimuli, even stress stimuli, contribute more cells to dominant unstressed clusters than to the less populated edgy clusters where stress dominates contribution. Thus, stressed cells tend towards the non-stress state. Unlike 250mM and 300mM sorbitol stress, 200mM sorbitol enables some growth and sufficient energy early central DGE more like both ND and NS. Lower stress enable ESC to simultaneous mount stress and developmental responses.

Bulk RNAseq showed marker gene sets for anterior visceral endoderm (AVE) being upregulated by ND and Sorbitol, which was not detected in any XEN clusters by scRNAseq (data not shown). We did not see any ND making XEN by scRNAseq analysis. This could mean that the results from bulk RNAseq data are predictive, and if the assay went longer in time, AVE markers would be found. Bulk RNAseq also indicated that XEN cells made is lowest in ND and increases dose-dependently. All cells will eventually make XEN, but ND is slow, and stresses are fast, which suggests that stress is causing premature differentiation by h is lineage imbalance.

In conclusion, we have shown here that less well-known hyperosmotic stress induces premature 2CEL XEN in all its three sub-lineages in a stress override of naïve stemness. The quality of the Sorbitol induction is like the RA control, but less powerful. The data suggests that stress stimuli produce more peripheral cells on the UMAP, and unstressed stimuli produce more central cells. Although each dose of stress contributes its highest number of cells to Clusters 0 and 1, which are each the largest cell clusters, stressed cells are proportionally lowest fraction in these clusters. In contrast, stress contributes high fractions, but relatively low numbers of cells to the peripheral (edgy) clusters. UMAP fingerprints and large marker sets support previous reports with small marker sets, that stress overrides stemness to produce premature enhances levels of 1^st^ lineage XEN cells. It also suggests that Stress fingerprints have the intrinsic capacity to map out the lineage and timing imbalances of different doses and types of stress, and to support interpretations of the dosimetric data from ESC HTS.

---

**Supplemental Figure 1.**
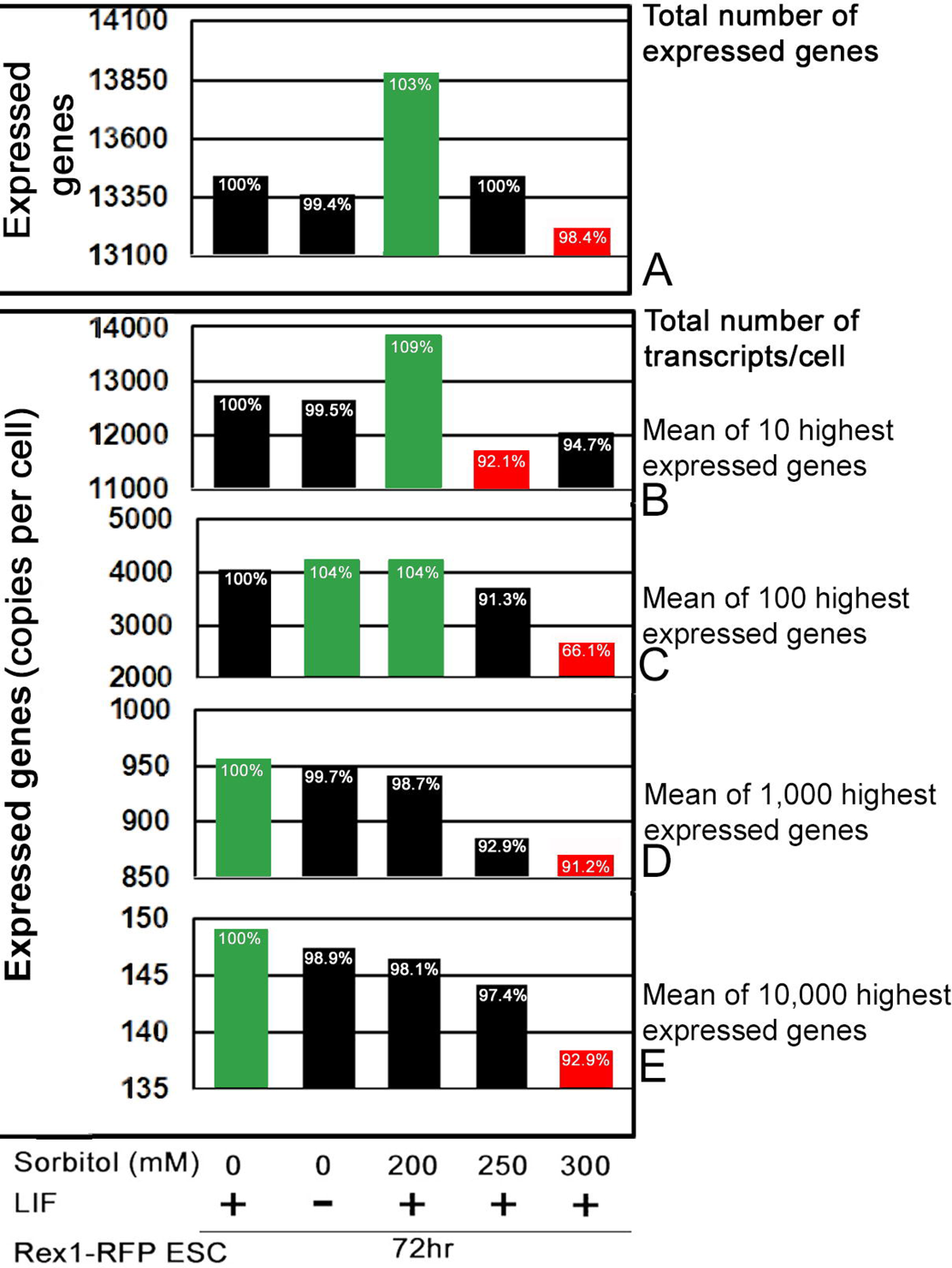
Rex1-RFP ESC were incubated with NS ND or 200-300mM sorbitol for 72hr and assay by bulk RNAseq with ND compared with NS for DGE during normal differentiation, and this was set to 100%, and stress forced changes compared to NS were then compared to ND/NS.

**Supplemental figure 2.**
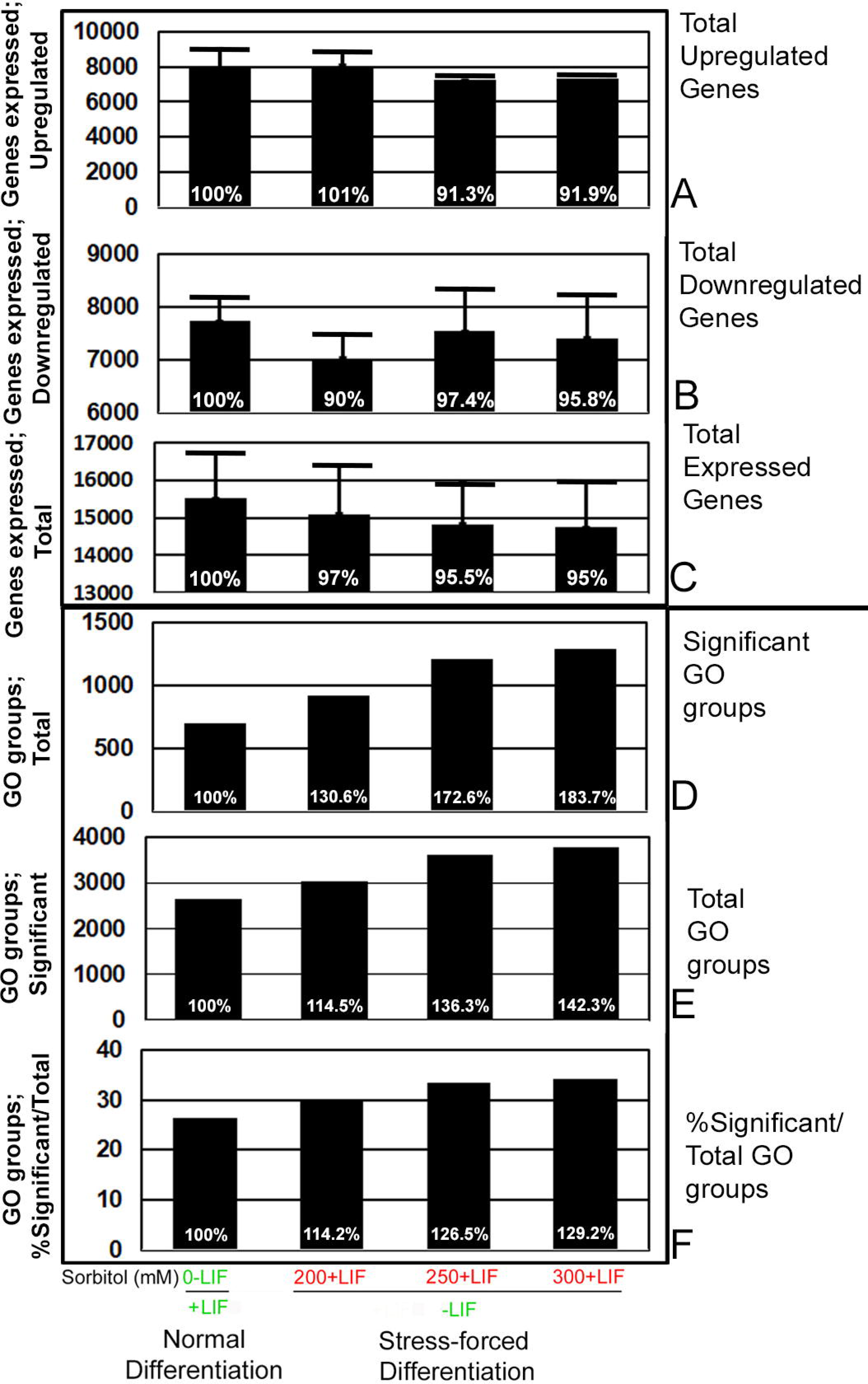
Stress decreases total genes expressed and transcripts/cell but increases total and significant GO groups. Rex1-RFP ESC were incubated with NS, ND, or 200-300mM sorbitol for 72hr and assayed by bulk RNAseq with ND compared with NS for DGE during normal differentiation, and this was set to 100%. The stress-forced changes compared to NS were then compared to ND/NS for single genes (**<u>A-C)</u>** and GO groups of genes (**<u>E-F</u>**).

**Supplemental figure 3.**
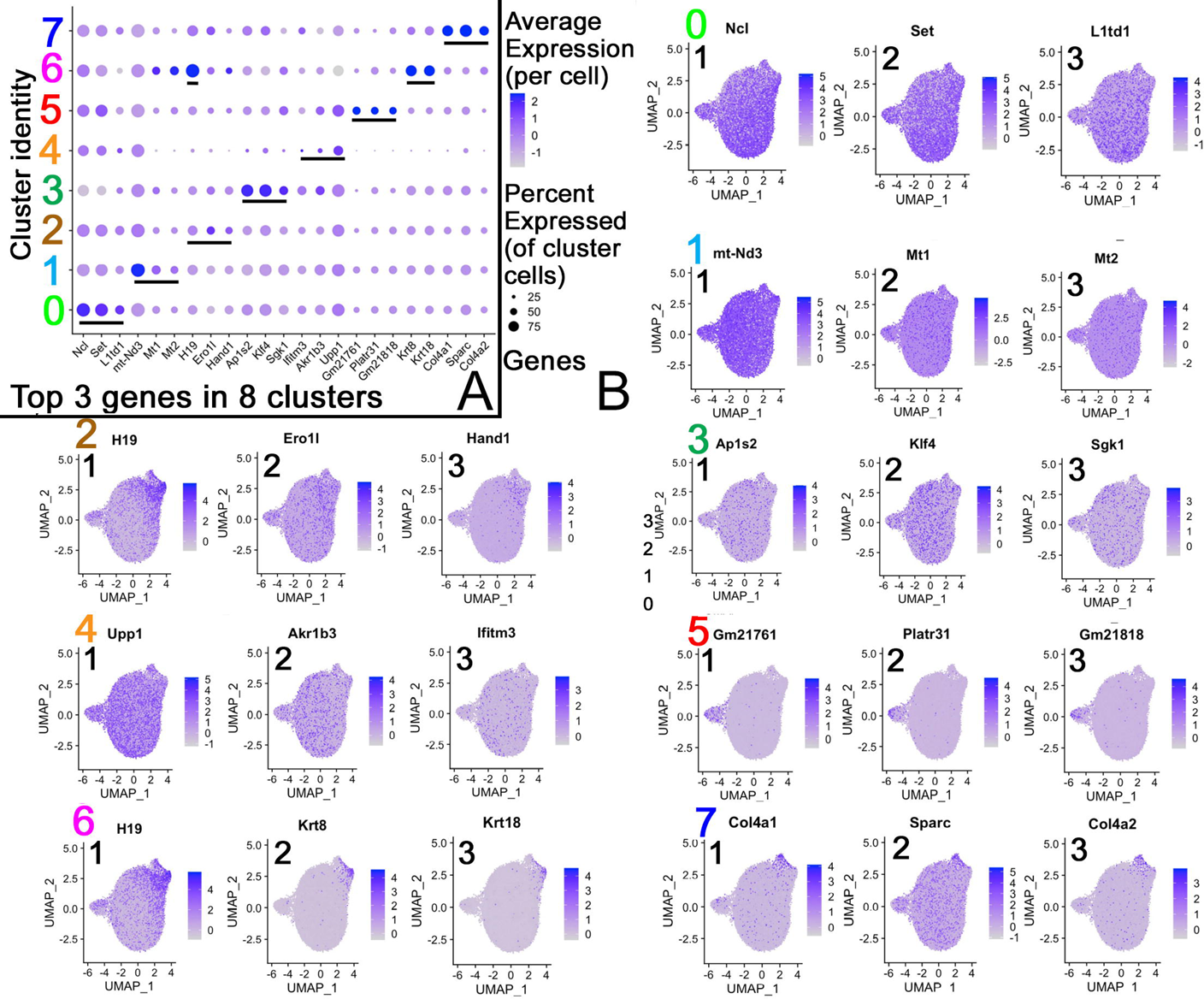
Top 3 bubble plots (**<u>A)</u>** and corresponding UMAPs for clusters 0-7 (**<u>B</u>**). Bubble plots for the top 3 genes for expression and subpopulation size initiate the definition of the 8 clusters by reading out the genes expressed at the highest levels and most cells/cluster.

**Supplemental figure 4.**
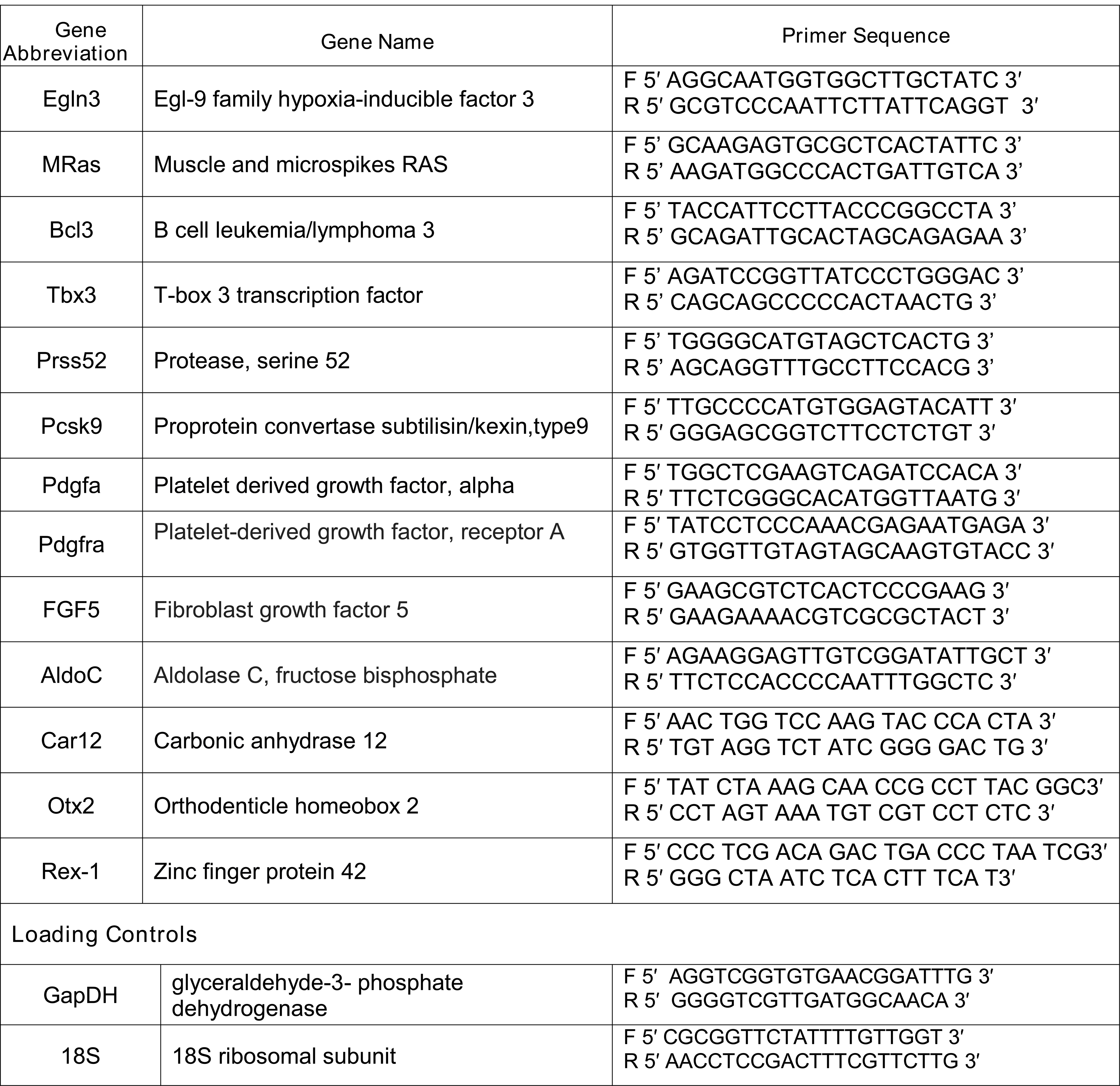
Primers for loading controls (Gapdh, 18S), developmental (Egl3, Otx2, FGF5, Tbx3, mRas, Pdgfa, Pdgfra, Rex1, Aldoc), and non-developmental genes (Car12, Bcl2, Prss52, Pcsk9) were assayed.

**Supplemental figure 5.**
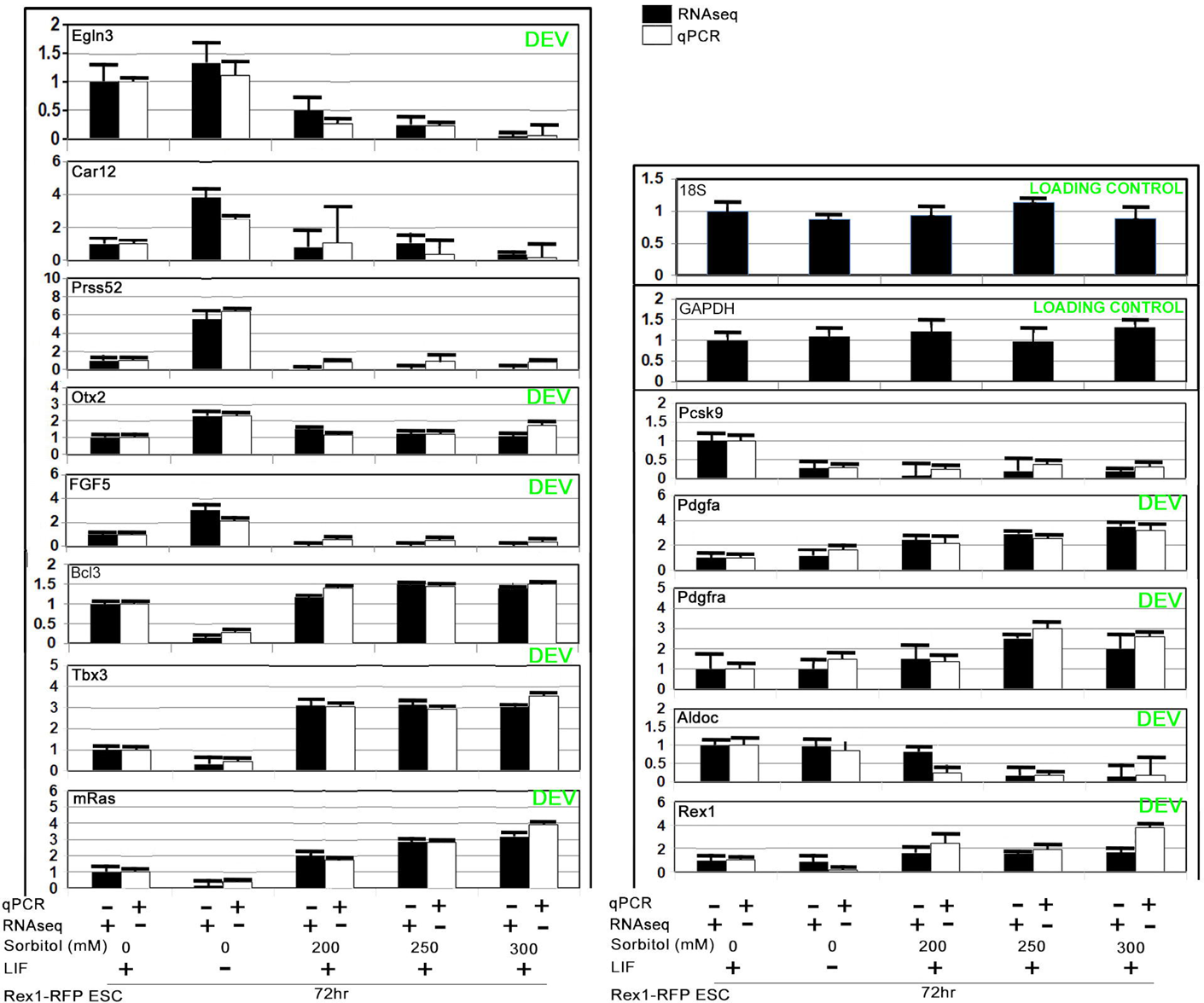
For the triplicate Rex1-RFP NS, ND, sorbitol 200, 250, and 300mM stimuli assayed by bulk RNAseq, triplicate measurements by the ddCt method of qPCR were assayed and graphed together in pairs. By t-test none of the genes were significantly different.

## Bibliography

Abdulhasan, M., Ruden, X., Rappolee, B., Dutta, S., Gurdziel, K., Ruden, D. M., Awonuga, A. O., Korzeniewski, S. J., Puscheck, E. E. and Rappolee, D. A. (2021a). Stress Decreases Host Viral Resistance and Increases Covid Susceptibility in Embryonic Stem Cells. Stem Cell Rev Rep 17, 2164–2177.

Abdulhasan, M. A., Ruden, X., Rappolee, B., Dutta, S., Gurdziel, K., Ruden, D. M., Awonuga, A. O., Korzeniewski, S. J., Puscheck, E. E. and Rappolee, D. A. (2021b). Stress decreases host viral resistance and increases Covid susceptibility in embryonic stem cells. Stem Cell Reviews and Reports 17, 2164–2177.

Anders, S., Pyl, P. T. and Huber, W. (2015). HTSeq--a Python framework to work with high-throughput sequencing data. Bioinformatics 31, 166–169.

Andrews, S. (2010). FastQC A Quality Control Tool for High Throughput Sequence Data.

Awonuga, A. O., Zhong, W., Abdallah, M. E., Slater, J. A., Zhou, S. C., Xie, Y. F., Puscheck, E. E. and Rappolee, D. A. (2011). Eomesodermin, HAND1, and CSH1 proteins are induced by cellular stress in a stress-activated protein kinase-dependent manner. Mol Reprod Dev 78, 519-528.

Betto, R. M., Diamante, L., Perrera, V., Audano, M., Rapelli, S., Lauria, A., Incarnato, D., Arboit, M., Pedretti, S., Rigoni, G., et al. (2021). Metabolic control of DNA methylation in naive pluripotent cells. Nat Genet 53, 215–229.

Boroviak, T., Loos, R., Lombard, P., Okahara, J., Behr, R., Sasaki, E., Nichols, J., Smith, A. and Bertone, P. (2015). Lineage-Specific Profiling Delineates the Emergence and Progression of Naive Pluripotency in Mammalian Embryogenesis. Dev Cell 35, 366–382.

Boroviak, T., Stirparo, G. G., Dietmann, S., Hernando-Herraez, I., Mohammed, H., Reik, W., Smith, A., Sasaki, E., Nichols, J. and Bertone, P. (2018). Single cell transcriptome analysis of human, marmoset and mouse embryos reveals common and divergent features of preimplantation development. Development 145.

Chan, M. M., Smith, Z. D., Grosswendt, S., Kretzmer, H., Norman, T. M., Adamson, B., Jost, M., Quinn, J. J., Yang, D., Jones, M. G., et al. (2019). Molecular recording of mammalian embryogenesis. Nature 570, 77–82.

Coronado, D., Godet, M., Bourillot, P. Y., Tapponnier, Y., Bernat, A., Petit, M., Afanassieff, M., Markossian, S., Malashicheva, A., Iacone, R., et al. (2013). A short G1 phase is an intrinsic determinant of naive embryonic stem cell pluripotency. Stem Cell Res 10, 118–131.

Despin-Guitard, E., Quenec’Hdu, R., Nahaboo, W., Schwarz, N., Leube, R. E., Chazaud, C. and Migeotte, I. (2022). Regionally specific levels and patterns of keratin 8 expression in the mouse embryo visceral endoderm emerge upon anterior-posterior axis determination. Front Cell Dev Biol 10, 1037041.

Detti, L., Francillon, L., Christiansen, M. E., Peregrin-Alvarez, I., Goedecke, P. J., Bursac, Z. and Roman, R. A. (2020a). Early pregnancy ultrasound measurements and prediction of first trimester pregnancy loss: A logistic model. Sci Rep 10, 1545.

Detti, L., Roman, R. A., Goedecke, P. J., Christiansen, M. E., Peregrin-Alvarez, I., Ikwuezunma, G. and Francillon, L. (2020b). Pilot study establishing a nomogram of yolk sac growth during the first trimester of pregnancy. J Obstet Gynaecol Res 46, 223–228.

Dobin, A., Davis, C. A., Schlesinger, F., Drenkow, J., Zaleski, C., Jha, S., Batut, P., Chaisson, M. and Gingeras, T. R. (2013). STAR: ultrafast universal RNA-seq aligner. Bioinformatics 29, 15–21.

Dong, C., Beltcheva, M., Gontarz, P., Zhang, B., Popli, P., Fischer, L. A., Khan, S. A., Park, K. M., Yoon, E. J., Xing, X., et al. (2020). Derivation of trophoblast stem cells from naive human pluripotent stem cells. Elife 9.

Dong, C. and Theunissen, T. W. (2022). Generating Trophoblast Stem Cells from Human Naive Pluripotent Stem Cells. Methods Mol Biol 2416, 91–104.

Emani, M. R., Narva, E., Stubb, A., Chakroborty, D., Viitala, M., Rokka, A., Rahkonen, N., Moulder, R., Denessiouk, K., Trokovic, R., et al. (2015). The L1TD1 protein interactome reveals the importance of post-transcriptional regulation in human pluripotency. Stem Cell Reports 4, 519–528.

Frum, T., Halbisen, M. A., Wang, C., Amiri, H., Robson, P. and Ralston, A. (2013). Oct4 cell-autonomously promotes primitive endoderm development in the mouse blastocyst. Dev Cell 25, 610–622.

Han, X., Wang, R., Zhou, Y., Fei, L., Sun, H., Lai, S., Saadatpour, A., Zhou, Z., Chen, H., Ye, F., et al. (2018). Mapping the Mouse Cell Atlas by Microwell-Seq. Cell 173, 1307.

Hao, Y., Hao, S., Andersen-Nissen, E., Mauck, W. M3rd., Zheng, S., Butler, A., Lee, M. J., Wilk, A. J., Darby, C., Zager, M., et al. (2021a). Integrated analysis of multimodal single-cell data. Cell 184, 3573-3587.e3529.

Hao, Y., Hao, S., Andersen-Nissen, E., Mauck, W. M3rd., Zheng, S., Butler, A., Lee, M. J., Wilk, A. J., Darby, C., Zager, M., et al. (2021b). Integrated analysis of multimodal single-cell data. Cell 184, 3573-3587 e3529.

Hu, Z., Tan, D. E. K., Chia, G., Tan, H., Leong, H. F., Chen, B. J., Lau, M. S., Tan, K. Y. S., Bi, X., Yang, D., et al. (2020). Maternal factor NELFA drives a 2C-like state in mouse embryonic stem cells. Nat Cell Biol 22, 175–186.

Huang, F.-J., Hsu, Y.-C., Kang, H.-Y., Chang, S.-Y., Hsuuw, Y.-D. and Huang, K.-E. (2005a). Effects of retinoic acid on the inner cell mass in mouse blastocysts. Fertility and Sterility 83, 238–242.

Huang, F.-J., Hsuuw, Y.-D., Lan, K.-C., Kang, H.-Y., Chang, S.-Y., Hsu, Y.-C. and Huang, K.-E. (2005b). Adverse effects of retinoic acid on embryo development and the selective expression of retinoic acid receptors in mouse blastocysts. Human Reproduction 21, 202–209.

Huang, F. J., Shen, C. C., Chang, S. Y., Wu, T. C. J. and Hsuuw, Y. D. (2003). Retinoic acid decreases the viability of mouse blastocysts in vitro. Human Reproduction 18, 130–136.

Huang, W., Sherman, B. T. and Lempicki, R. A. (2009). Systematic and integrative analysis of large gene lists using DAVID bioinformatics resources. Nature protocols 4, 44–57.

Illingworth, R. S., Holzenspies, J. J., Roske, F. V., Bickmore, W. A. and Brickman, J. M. (2016). Polycomb enables primitive endoderm lineage priming in embryonic stem cells. Elife 5.

Iturbide, A., Ruiz Tejada Segura, M. L., Noll, C., Schorpp, K., Rothenaigner, I., Ruiz-Morales, E. R., Lubatti, G., Agami, A., Hadian, K., Scialdone, A., et al. (2021). Retinoic acid signaling is critical during the totipotency window in early mammalian development. Nat Struct Mol Biol 28, 521–532.

Iturbide, A., Ruiz Tejada Segura, M. L., Noll, C., Schorpp, K., Rothenaigner, I., Ruiz-Morales, E. R., Lubatti, G., Agami, A., Hadian, K., Scialdone, A., et al. (2022). Author Correction: Retinoic acid signaling is critical during the totipotency window in early mammalian development. Nat Struct Mol Biol 29, 282.

Iwabuchi, K. A., Yamakawa, T., Sato, Y., Ichisaka, T., Takahashi, K., Okita, K. and Yamanaka, S. (2011). ECAT11/L1td1 is enriched in ESCs and rapidly activated during iPSC generation, but it is dispensable for the maintenance and induction of pluripotency. PLoS One 6, e20461.

Judson, R., Houck, K., Martin, M., Richard, A. M., Knudsen, T. B., Shah, I., Little, S., Wambaugh, J., Woodrow Setzer, R., Kothiya, P., et al. (2016). Editor’s Highlight: Analysis of the Effects of Cell Stress and Cytotoxicity on In Vitro Assay Activity Across a Diverse Chemical and Assay Space. Toxicol Sci 152, 323-339.

Kalkan, T., Olova, N., Roode, M., Mulas, C., Lee, H. J., Nett, I., Marks, H., Walker, R., Stunnenberg, H. G., Lilley, K. S., et al. (2017). Tracking the embryonic stem cell transition from ground state pluripotency. Development 144, 1221–1234.

Kalkan, T. and Smith, A. (2014). Mapping the route from naive pluripotency to lineage specification. Philos Trans R Soc Lond B Biol Sci 369.

Khan, S. A., Park, K. M., Fischer, L. A., Dong, C., Lungjangwa, T., Jimenez, M., Casalena, D., Chew, B., Dietmann, S., Auld, D. S., et al. (2021). Probing the signaling requirements for naive human pluripotency by high-throughput chemical screening. Cell Rep 35, 109233.

Kinoshita, M., Li, M. A., Barber, M., Mansfield, W., Dietmann, S. and Smith, A. (2021). Disabling de novo DNA methylation in embryonic stem cells allows an illegitimate fate trajectory. Proc Natl Acad Sci U S A 118.

Knoetgen, H., Teichmann, U. and Kessel, M. (1999). Head-organizing activities of endodermal tissues in vertebrates. Cell Mol Biol (Noisy-le-grand) 45, 481–492.

Kondoh, H. (2018). Roles of ZIC2 in Regulation of Pluripotent Stem Cells. Adv Exp Med Biol 1046, 339–351.

Li, L., Arman, E., Ekblom, P., Edgar, D., Murray, P. and Lonai, P. (2004). Distinct GATA6- and laminin-dependent mechanisms regulate endodermal and ectodermal embryonic stem cell fates. Development 131, 5277–5286.

Li, Q., Gomez-Lopez, N., Drewlo, S., Sanchez-Rodriguez, E., Dai, J., Puscheck, E. E. and Rappolee, D. A. (2016a). Development and Validation of a Rex1-RFP Potency Activity Reporter Assay That Quantifies Stress-Forced Potency Loss in Mouse Embryonic Stem Cells. Stem Cells Dev 25, 320–328.

Li, Q., Louden, E., Zhou, J., Drewlo, S., Dai, J., Puscheck, E. E., Chen, K. and Rappolee, D. A. (2019). Stress Forces First Lineage Differentiation of Mouse Embryonic Stem Cells; Validation of a High-Throughput Screen for Toxicant Stress. Stem Cells Dev 28, 101–113.

Li, Q., Yang, Y., Louden, E., Puscheck, E. and Rappolee, D. (2016b). High throughput screens for embryonic stem cells; stress-forced potency-stemness loss enables toxicological assays. In Methods In Toxicology and Pharmacology (ed. A. Faqi): Springer.

Livak, K. J. and Schmittgen, T. D. (2001). Analysis of relative gene expression data using real-time quantitative PCR and the 2(-Delta Delta C(T)) Method. Methods 25, 402–408.

Madabhushi, M. and Lacy, E. (2011). Anterior visceral endoderm directs ventral morphogenesis and placement of head and heart via BMP2 expression. Dev Cell 21, 907–919.

Marks, H., Kalkan, T., Menafra, R., Denissov, S., Jones, K., Hofemeister, H., Nichols, J., Kranz, A., Stewart, A. F., Smith, A., et al. (2012). The transcriptional and epigenomic foundations of ground state pluripotency. Cell 149, 590–604.

Martinez-Ceballos, E., Chambon, P. and Gudas, L. J. (2005). Differences in gene expression between wild type and Hoxa1 knockout embryonic stem cells after retinoic acid treatment or leukemia inhibitory factor (LIF) removal. J Biol Chem 280, 16484–16498.

Masui, S., Ohtsuka, S., Yagi, R., Takahashi, K., Ko, M. S. and Niwa, H. (2008). Rex1/Zfp42 is dispensable for pluripotency in mouse ES cells. BMC Dev Biol 8, 45.

Mohammed, H., Hernando-Herraez, I., Savino, A., Scialdone, A., Macaulay, I., Mulas, C., Chandra, T., Voet, T., Dean, W., Nichols, J., et al. (2017). Single-Cell Landscape of Transcriptional Heterogeneity and Cell Fate Decisions during Mouse Early Gastrulation. Cell Rep 20, 1215–1228.

Murray, P. and Edgar, D. (2001). Regulation of the differentiation and behaviour of extra-embryonic endodermal cells by basement membranes. J Cell Sci 114, 931–939.

Neagu, A., van Genderen, E., Escudero, I., Verwegen, L., Kurek, D., Lehmann, J., Stel, J., Dirks, R. A. M., van Mierlo, G., Maas, A., et al. (2020). In vitro capture and characterization of embryonic rosette-stage pluripotency between naive and primed states. Nat Cell Biol 22, 534–545.

Niwa, H., Miyazaki, J.-i. and Smith, A. G. (2000). Quantitative expression of Oct-3/4 defines differentiation, dedifferentiation or self-renewal of ES cells. Nat Genet 24, 372–376.

Palmieri, S. L., Peter, W., Hess, H. and Scholer, H. R. (1994). Oct-4 transcription factor is differentially expressed in the mouse embryo during establishment of the first two extraembryonic cell lineages involved in implantation. Dev Biol 166, 259–267.

Percharde, M., Lin, C. J., Yin, Y., Guan, J., Peixoto, G. A., Bulut-Karslioglu, A., Biechele, S., Huang, B., Shen, X. and Ramalho-Santos, M. (2018). A LINE1-Nucleolin Partnership Regulates Early Development and ESC Identity. Cell 174, 391–405 e319.

Posfai, E., Schell, J. P., Janiszewski, A., Rovic, I., Murray, A., Bradshaw, B., Yamakawa, T., Pardon, T., El Bakkali, M., Talon, I., et al. (2021). Evaluating totipotency using criteria of increasing stringency. Nat Cell Biol 23, 49–60.

Puscheck, E. E., Awonuga, A. O., Yang, Y., Jiang, Z. and Rappolee, D. A. (2015). Molecular biology of the stress response in the early embryo and its stem cells. Adv Exp Med Biol 843, 77–128.

Puscheck, E. E., Ruden, X., Singh, A., Abdulhasan, M., Ruden, D. M., Awonuga, A. O. and Rappolee, D. A. (2022). Using high throughput screens to predict miscarriages with placental stem cells and long-term stress effects with embryonic stem cells. Birth Defects Res.

Rappolee, D. A. (1999). It’s not just baby’s babble/Babel: recent progress in understanding the language of early mammalian development: a minireview. Mol Reprod Dev 52, 234–240.

Rappolee, D. A., Xie, Y., Slater, J. A., Zhou, S. and Puscheck, E. E. (2012). Toxic stress prioritizes and imbalances stem cell differentiation: implications for new biomarkers and in vitro toxicology tests. Syst Biol Reprod Med 58, 33–40.

Robinson, M. D., McCarthy, D. J. and Smyth, G. K. (2010). edgeR: a Bioconductor package for differential expression analysis of digital gene expression data. Bioinformatics 26, 139–140.

Rodriguez-Terrones, D., Gaume, X., Ishiuchi, T., Weiss, A., Kopp, A., Kruse, K., Penning, A., Vaquerizas, J. M., Brino, L. and Torres-Padilla, M. E. (2018). A molecular roadmap for the emergence of early-embryonic-like cells in culture. Nat Genet 50, 106–119.

Rodriguez, A., Diez, C., Ikeda, S., Royo, L. J., Caamano, J. N., Alonso-Montes, C., Goyache, F., Alvarez, I., Facal, N. and Gomez, E. (2006). Retinoids during the in vitro transition from bovine morula to blastocyst. Hum Reprod 21, 2149–2157.

Ross, C. and Boroviak, T. E. (2020). Origin and function of the yolk sac in primate embryogenesis. Nature Communications 11, 3760.

Rutledge, J. C., Shourbaji, A. G., Hughes, L. A., Polifka, J. E., Cruz, Y. P., Bishop, J. B. and Generoso, W. M. (1994). Limb and lower-body duplications induced by retinoic acid in mice. Proc Natl Acad Sci U S A 91, 5436–5440.

Slater, J. A., Zhou, S., Puscheck, E. E. and Rappolee, D. A. (2014). Stress-induced enzyme activation primes murine embryonic stem cells to differentiate toward the first extraembryonic lineage. Stem Cells Dev 23, 3049–3064.

Stuart, T., Butler, A., Hoffman, P., Hafemeister, C., Papalexi, E., Mauck, W. M3rd., Hao, Y., Stoeckius, M., Smibert, P. and Satija, R. (2019). Comprehensive Integration of Single-Cell Data. Cell 177, 1888-1902 e1821.

Theunissen, T. W., Powell, B. E., Wang, H., Mitalipova, M., Faddah, D. A., Reddy, J., Fan, Z. P., Maetzel, D., Ganz, K., Shi, L., et al. (2014). Systematic identification of culture conditions for induction and maintenance of naive human pluripotency. Cell Stem Cell 15, 471–487.

Tokuzawa, Y., Kaiho, E., Maruyama, M., Takahashi, K., Mitsui, K., Maeda, M., Niwa, H. and Yamanaka, S. (2003). Fbx15 is a novel target of Oct3/4 but is dispensable for embryonic stem cell self-renewal and mouse development. Mol Cell Biol 23, 2699–2708.

Wang, H. Y., Kanungo, J. and Malbon, C. C. (2002). Expression of Galpha 13 (Q226L) induces P19 stem cells to primitive endoderm via MEKK1, 2, or 4. J Biol Chem 277, 3530-3536.

Wang, X., Xiang, Y., Yu, Y., Wang, R., Zhang, Y., Xu, Q., Sun, H., Zhao, Z.-A., Jiang, X., Wang, X., et al. (2021a). Formative pluripotent stem cells show features of epiblast cells poised for gastrulation. Cell Research 31, 526–541.

Wang, Y., Na, Q., Li, X., Tee, W. W., Wu, B. and Bao, S. (2021b). Retinoic acid induces NELFA-mediated 2C-like state of mouse embryonic stem cells associates with epigenetic modifications and metabolic processes in chemically defined media. Cell Prolif 54, e13049.

Wong, L. H., McGhie, J. D., Sim, M., Anderson, M. A., Ahn, S., Hannan, R. D., George, A. J., Morgan, K. A., Mann, J. R. and Choo, K. H. (2010). ATRX interacts with H3.3 in maintaining telomere structural integrity in pluripotent embryonic stem cells. Genome Res 20, 351–360.

Xie, Y., Awonuga, A., Liu, J., Rings, E., Puscheck, E. E. and Rappolee, D. A. (2013). Stress Induces AMPK-Dependent Loss of Potency Factors Id2 and Cdx2 in Early Embryos and Stem Cells. Stem Cells and Development 22, 1564–1575.

Xie, Y., Zhong, W., Wang, Y., Trostinskaia, A., Wang, F., Puscheck, E. E. and Rappolee, D. A. (2007). Using hyperosmolar stress to measure biologic and stress-activated protein kinase responses in preimplantation embryos. Mol Hum Reprod 13, 473–481.

Yang, S.-H., Kalkan, T., Morissroe, C., Marks, H., Stunnenberg, H., Smith, A. and Sharrocks, Andrew D. (2014). Otx2 and Oct4 Drive Early Enhancer Activation during Embryonic Stem Cell Transition from Naive Pluripotency. Cell Reports 7, 1968–1981.

Yang, Y., Arenas-Hernandez, M., Gomez-Lopez, N., Dai, J., Parker, G. C., Puscheck, E. E. and Rappolee, D. A. (2016). Hypoxic Stress Forces Irreversible Differentiation of a Majority of Mouse Trophoblast Stem Cells Despite FGF4. Biol Reprod 95, 110.

Zheng, G. X., Terry, J. M., Belgrader, P., Ryvkin, P., Bent, Z. W., Wilson, R., Ziraldo, S. B., Wheeler, T. D., McDermott, G. P., Zhu, J., et al. (2017). Massively parallel digital transcriptional profiling of single cells. Nat Commun 8, 14049.

Zhong, W., Xie, Y., Abdallah, M., Awonuga, A. O., Slater, J. A., Sipahi, L., Puscheck, E. E. and Rappolee, D. A. (2010). Cellular stress causes reversible, PRKAA1/2-, and proteasome-dependent ID2 protein loss in trophoblast stem cells. Reproduction 140, 921-930.

Zhong, W., Xie, Y., Wang, Y., Lewis, J., Trostinskaia, A., Wang, F., Puscheck, E. E. and Rappolee, D. A. (2007). Use of hyperosmolar stress to measure stress-activated protein kinase activation and function in human HTR cells and mouse trophoblast stem cells. Reprod Sci 14, 534–547.

Zhou, X. and Anderson, K. V. (2010). Development of head organizer of the mouse embryo depends on a high level of mitochondrial metabolism. Dev Biol 344, 185–195.

Zhuang, Y. and Gudas, L. J. (2008). Overexpression of COUP-TF1 in murine embryonic stem cells reduces retinoic acid-associated growth arrest and increases extraembryonic endoderm gene expression. Differentiation 76, 760–771.

